# Long-range cortical synchronization supports abrupt visual learning

**DOI:** 10.1101/2021.08.03.454994

**Authors:** Bennett A. Csorba, Matthew R. Krause, Theodoros P. Zanos, Christopher C. Pack

**Affiliations:** Montreal Neurological Institute, McGill University, Montreal, QC H3A 2B4, Canada; Feinstein Institute for Medical Research, Manhasset, NY 11030, USA

**Keywords:** Learning, vision, neuronal oscillations, inferotemporal cortex, prefrontal cortex, neural synchronization

## Abstract

Visual plasticity declines sharply after the critical period, yet we easily learn to recognize new faces and places even as adults. Such learning is often characterized by a “moment of insight”, an abrupt and dramatic improvement in recognition. The mechanisms that support abrupt learning are unknown, but one hypothesis is that they involve changes in synchronization between brain regions. To test this hypothesis, we used a behavioral task in which non-human primates rapidly learned to recognize novel images and to associate them with specific responses. Simultaneous recordings from inferotemporal and prefrontal cortices revealed a transient synchronization of neural activity between these areas that peaked around the moment of insight. Synchronization was strongest between inferotemporal sites that encoded images and reward-sensitive prefrontal sites. Moreover, its magnitude intensified gradually over image exposures, suggesting that abrupt learning culminates from the search for informative signals within a circuit linking sensory information to task demands.

## INTRODUCTION

In adults, visual learning often requires prolonged training. Even for simple tasks, such as discriminating the orientation of a line, behavioral changes often emerge after days or weeks of practice^1^. For more complex tasks, such as the detection of anomalies in medical images, efficient performance requires months or years of training^2, 3^. Neurophysiological studies have similarly revealed that the adult visual cortex often changes very slowly, if at all, in response to experience^4, 5^.

At the same time, it is clear that adults are capable of rapid visual learning under the right circumstances^6, 7^. Indeed, learning can even occur following a single exposure to a stimulus^8^, or *abruptly,* after a series of unsuccessful attempts at a task^6, 7^. This latter kind of learning, which Hebb referred to as “insight”^9^, has frequently been observed in freely behaving animals attempting to obtain rewards in unfamiliar settings. These observations pose a challenge for modern theories of learning that rely on gradual synaptic changes following many presentations of the same stimulus^10^.

From an ethological perspective, abrupt learning would seem to be necessary for survival in natural visual environments, which typically do not afford the opportunity for hundreds of exposures to novel stimuli. At the same time, such plasticity must be engaged selectively, to prevent newly learned stimuli from overwhelming existing representations. One solution to this “stability-plasticity” dilemma is therefore to impose a gating mechanism, whereby abrupt visual learning occurs only when specific subgroups of neurons are active together^9,11^. For visual learning, the relevant subgroups might be those that encode the relevant stimuli and those that encode the demands of a given task or context^12^.

At a neural level, these operations can be implemented through oscillatory *synchronization,* which has been shown to support the rapid formation of new memories^13–15^, as well as long-range communication more generally^16^. It has also been implicated in functions that are important for slower forms of learning, such as attention^1^ and reward sensitivity^14^. Critically, oscillatory synchronization can change flexibly on short time-scales, so as to link different brain regions that contain different types of task-relevant information^11^. However, the role of this kind of synchronization in abrupt visual learning is unknown.

Here, we have tested the hypothesis that oscillatory synchronization facilitates rapid visual learning, using multi-site neural recordings in non-human primates. The animals were trained to perform a naturalistic “foraging” task, in which they learned to recognize a visual image and to associate it with a rewarded location. Learning in this task was abrupt, with most sessions being characterized by large performance improvements over the course of a few trials. At the same time, we recorded from the prefrontal and inferotemporal cortices, which are known to interact during visual learning and perception^17–19^.

Around the “moment of insight”, we found a transient increase in synchronization between neural activity in these two cortical areas. This increase in synchronization was strongest between prefrontal sites that encoded rewards and inferotemporal sites that discriminated between the relevant visual stimuli. In contrast, we did not find local changes in neural firing or oscillatory power that correlated strongly with learning. These results therefore suggest that rapid learning relies on temporal synchronization between cortical sites that connect relevant stimuli with task outcomes.

## RESULTS

### Abrupt visual learning in a naturalistic behavioral paradigm

To probe the mechanisms of abrupt learning, we used the “oculomotor foraging” task shown in Figure 1A^20, 21^. On each trial, animals freely explored a natural scene until their gaze landed within an unmarked *reward zone* (RZ). A fixation within the RZ triggered the release of a few drops of juice and ended the trial. Both the starting eye position and the precise location of the RZ varied from trial to trial (see Methods), so that animals could not perform the task by simply associating a fixed saccade vector with each image (Figure S1, panels G-L). Instead, they had to learn the spatiotopic location of the RZ within the image.

**Figure 1.**
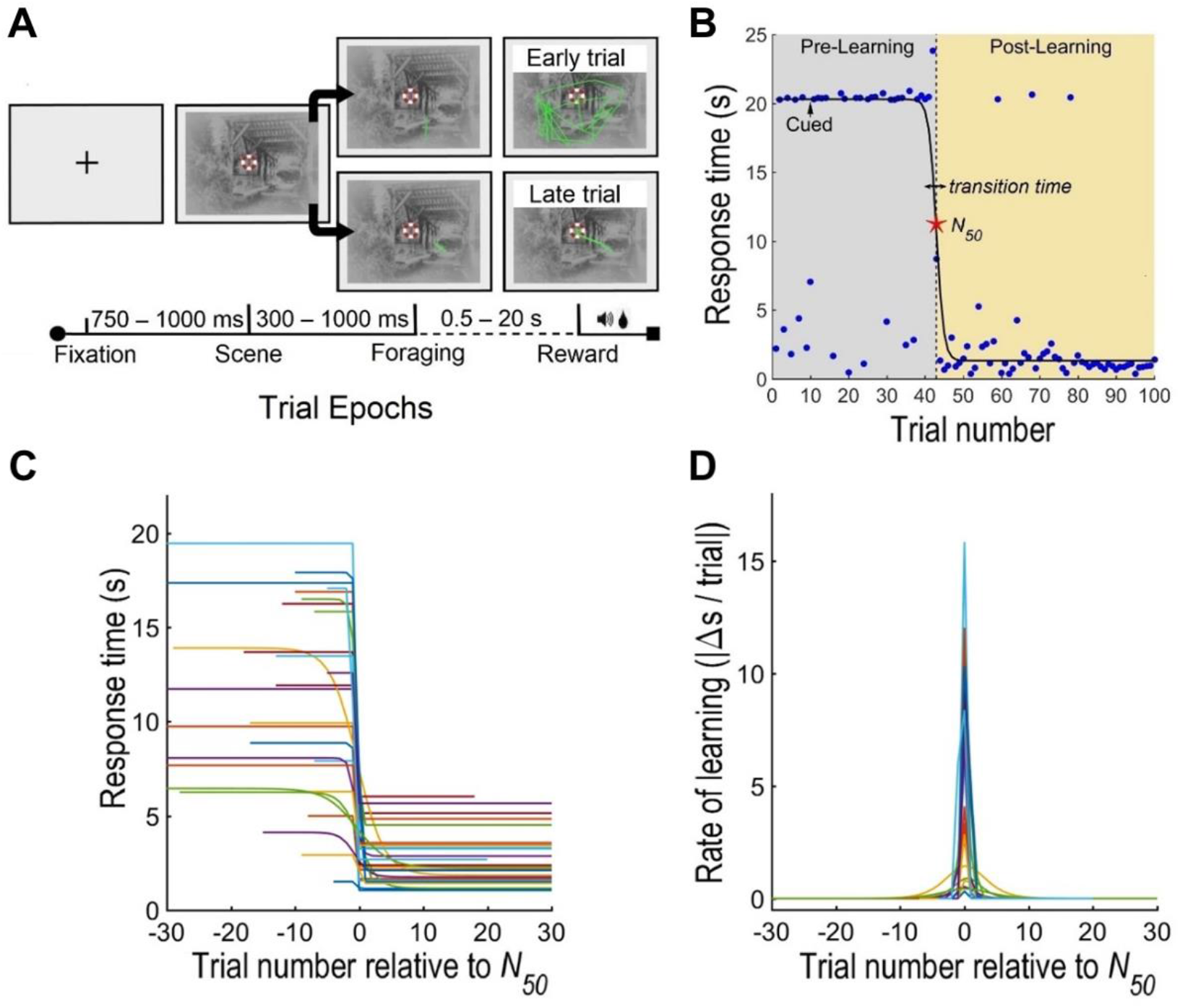
Task diagram and behavior. (A) An oculomotor “foraging” task was used to study visual learning. Each trial began with the onset of a fixation cross. After fixation was acquired, a natural image appeared, and animals were then free to explore the image in search of a reward zone (marked here with a red-white ring, but not visible to the subject). Fixation on the reward zone led to the release of the reward and the end of the trial. If the reward zone was not found after 15 s (Monkey F) or 20 s (Monkey M), a cue indicated the rewarded location. Each trial was therefore characterized by a Scene epoch, a Foraging epoch, and a Reward epoch. Eye traces (green) are shown for an example early trial (top), when the animal failed to find the reward zone after 20 s, and a late trial (bottom), after learning was complete. Two images were shown in an interleaved manner on every session. (B) Behavioral performance for an example image. Blue dots indicate the time required to find the target on a single trial, and the black line indicates the fit of a sigmoid function to the data for all trials. The red star indicates the *N_50_* trial, estimated from the sigmoid fit as the time required for performance to improve by 50% from its pre-learning to its post-learning state. (C) Sigmoid fits for all images for Monkey M, aligned on the *N_50_* value for each image. Fits for Monkey F are included in Figure S1E. (D) The rate of learning, defined as the performance change as a function of trial, for all images for Monkey M, aligned on the *N_50_* value for each image. Data for Monkey F are included in Figure S1 (panel F).

Within each session, animals were exposed to two different images, each with its own unique RZ, in a randomly interleaved fashion. Visual learning was therefore an important component of the task, as both the images were initially unfamiliar to the animals. Through repeated presentations of each image, the animals learned to recognize them and to associate them with the corresponding RZs.

Figure 1B shows the progression of learning within a single example session, considering only one of the two images presented. Each point shows the *response time*^21^, defined as the time it took the animal to find the RZ on each trial. Although the animal occasionally found the RZ quickly, typical response times were initially very high. This was not due to a lack of engagement, as the animal actively searched the image, making on average 41.5 saccades on each trial (Figure S1B). To maintain engagement during the early pre-learning phase, these trials timed out after 15 or 20 s, at which point a cue was shown to provide a hint about the correct location (Figure 1A, top).

On the 43^rd^ trial for this example image, a “moment of insight” occurred, and the typical response time abruptly dropped from 20 s to ~1 s. Response times then remained low on most subsequent trials, suggesting that the animal had successfully learned to recognize this image and to associate it with the corresponding RZ. Overall task engagement appeared to be similar before and after learning, as the animals continued to generate saccades at similar rates (Figure S1B; 3.69 ± 0.27 vs. 3.29 ± 0.21 saccades/s). However, after learning the saccades were generally directed toward the RZ, resulting in trials of shorter duration.

To quantify the dynamics of learning, we fit the sequence of response times for each image to a sigmoid function (solid line in Figure 1B). From the sigmoid fits, we extracted two quantities (see *Methods).* The first was *N_50_,* (red star in Figure 1B), defined as the trial at which response time decreased by half from its initial value; we used this value as an objective marker of the trial around which learning was centered. Secondly, the *transition time* (arrow in Figure 1B) was defined as the number of trials required for performance to go from 75% to 25% of the maximum response time indicated by the sigmoid fit. For the data shown in Figure 1 (panel B), the transition time was 2 trials, which lasted 10.1 seconds in total, indicating that learning was indeed abrupt.

In total, we obtained behavioral and electrophysiological data from sessions involving 50 different images (37 from monkey M and 13 from monkey F). As in the example, the behavioral data were well fit by a sigmoid function (median r^2^ = 0.76), with the fit quality not differing significantly between the two animals (one-way ANOVA, p = 0.59). From these fits, we estimate that the median value of *N_50_* was 18 trials, and the mean transition time was 1.58 trials *(SE* = 0.15 trials). Basic eye movement metrics, including saccade rate, microsaccade rate, and latency to the first saccade on each trial did not vary significantly with learning or between subjects (two-way ANOVAs, *p* > 0.05).

Figures 1C and 1D summarize the progression and rate of learning for all images for an individual animal (Monkey M). As in the example session, performance changes were confined to a few trials around the *N_50_* trial for each image. Similar results for the second animal are shown in Figure S1E-F. Overall, these results show that learning had a pronounced effect on task performance; it happened rapidly; and it usually followed a prolonged period of unsuccessful task performance. These characteristics are typical of abrupt learning^6, 9^.

We also verified that this learning was *persistent,* by retesting the animals on a subset of the images, many days after the first exposure. On average, response times during the first 10 trials for each image decreased by 1.90 s from the first session to the retest session, and the *N_50_* value decreased by 12.6 trials (two-way ANOVAs, by session, *p* < 0.05, for animal or session x animal interaction, *p* > 0.05). Even before *N_50_*, animals consistently made fewer saccades during recall sessions than the initial sessions (two-way ANOVA, *p* < 0.05), indicating that they retained a memory of the images. Thus, our foraging paradigm led to abrupt learning that was quite durable, as has been reported for other abrupt learning tasks^22^ and for learning in the wild^23^.

### Cortical dynamics of abrupt learning

To assess the cortical basis of abrupt learning, we recorded simultaneously from 96-channel electrode arrays in the inferotemporal (IT) cortex and the prefrontal cortex (PFC) (Figure 2). Area IT is known to be important for recognizing complex images^24^, while PFC plays a role in learning arbitrary associations^25^. Lesion studies have suggested that the connections between these two areas are especially important for learning associations between images and spatial locations^19, 26^, but the physiological nature of this interaction is unknown^27^. We therefore examined the functional interactions between sites in IT and PFC, to determine how they evolve with learning.

**Figure 2.**
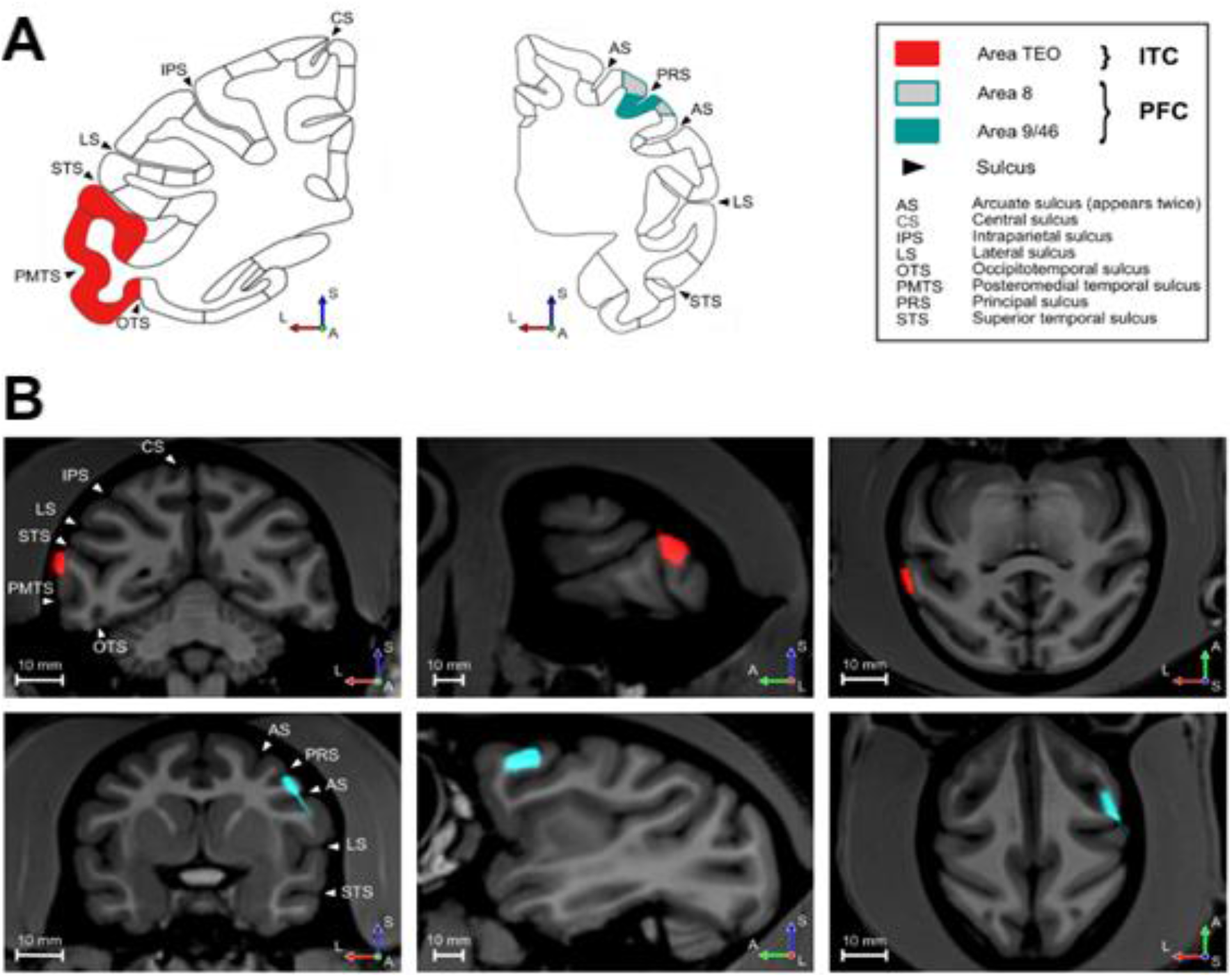
Electrode Array Locations (Monkey F). Images are shown in neurological convention (“left is left”). L: Left, A: Anterior, S: Superior. (A) Coronal sections from the Markov et al. CoreNets atlas^30^ showing the brain areas targeted for recording. The sulcal landmarks indicated were used to identify the same regions in each animal’s MRI. Due to its curved shape, the arcuate sulcus (AS) appears twice in the coronal PFC sections. (B) Combined MRI/CT images showing the positions of the IT (top row) and PFC (bottom) arrays in coronal (left), sagittal (center), and axial (right) planes. The preoperative T1-weighted MRI was co-registered with a postoperative CT scan (red, cyan) to verify the array locations. To emphasize the neuroanatomy, the skull and artifacts from other implants have been digitally removed from the CT scan.

To focus our analysis, we defined three epochs during each behavioral trial. As shown in Figure 1A, these are the “Scene Onset” epoch, when the image first appeared; the “Foraging” epoch, when search was initiated; and the “Reward” epoch, when the reward was dispensed. These were chosen because retinal stimulation was largely identical within each epoch throughout the progression of learning (see Methods). To characterize learning-related changes in synchronization between PFC and IT, we examined the *oscillatory coherence* between the local field potential (LFP) signals in each area, a commonly used metric that indexes transient changes in corticocortical communication during learning^28, 29^ (see Methods).

To illustrate the general pattern of results, Figure 3A shows the magnitude of synchronization between the two areas, averaged across electrodes and images for both animals. Here, we have grouped trials into a pre-learning phase (before *N_50_*, top row), a post-learning phase (after *N_50_*, bottom row) and those around the moment of abrupt learning (*N_50_*, middle row). The left column, corresponding to the Scene Onset epoch, shows that the appearance of a visual image generally triggered an increase in low-frequency synchronization starting around 200 ms later. This sensory response was consistent throughout the progression of learning (two-way ANOVA, either learning phase or by animal or the interaction, *p* > 0.05).

**Figure 3.**
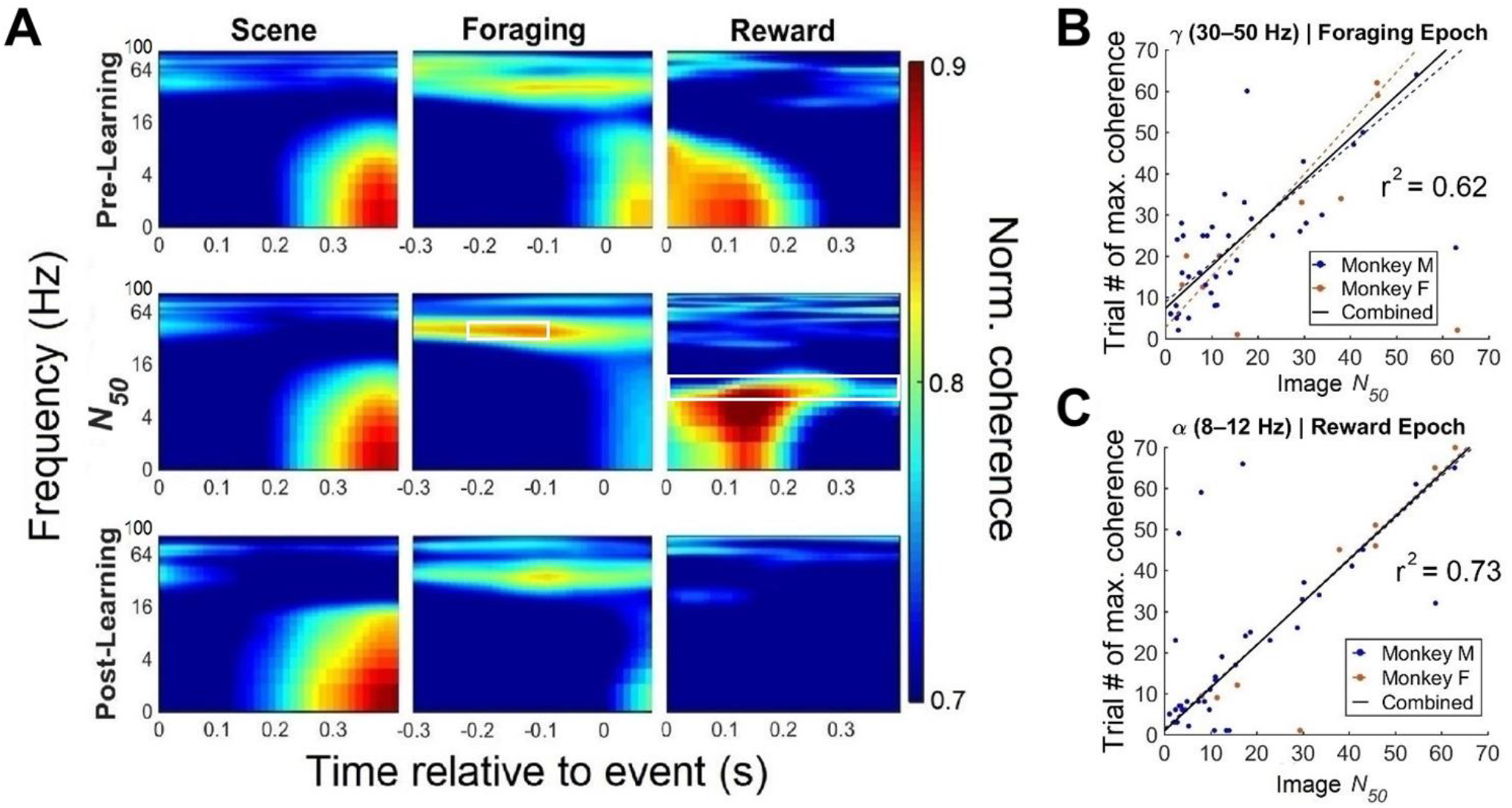
Inferotemporal-prefrontal oscillatory synchronization reflects learning state. (A) Synchronization of LFP signals between IT and PFC are shown for the three trial epochs (Scene, Foraging, and Reward) at different stages of learning (Pre-Learning, *N_50_*, and Post-Learning). Each panel contains a time-frequency plot covering frequencies in the range 1 - 100 Hz and a period of 400 ms around the event that defines each epoch. For the Scene and Reward epochs, t = 0 is the onset of the image and reward respectively. For the Foraging epoch, t = 0 is the onset of the first saccade within a trial. The strength of synchronization is normalized within each frequency and averaged across individual sessions. Time-frequency regions of interest selected from these preliminary data are indicated in white for the Foraging epoch (middle column, middle row) and the Reward epoch (right column, middle row). Around the moment of learning, there was a notable increase in gamma (30-50 Hz) synchronization in the Foraging epoch (middle column, middle row, two-sided permutation t-tests, *p* < 0.05), as well as increased alpha band (8-12 Hz) synchronization during the Reward epoch (right column, middle row, two-sided permutation t-tests, *p* < 0.05) relative to pre-learning activity. (B) Trial number of peak gamma synchronization (30-50 Hz) in the Foraging epoch is plotted against the *N_50_* value from the sigmoid fit for the corresponding behavioral data, for each monkey separately (red and blue dots), along with best-fitting regression lines for each animal (red and blue dashed line). The black line indicates the best-fitting regression line for all data combined. (C) As in (B), but for alpha synchronization during the Reward epoch.

In contrast, the onset of the Foraging epoch (middle column) was accompanied by an increase in gamma band (30–50 Hz) synchronization, which was most prominent around the *N_50_* trial, when learning was most evident (middle row). A Granger-Geweke causality analysis revealed that the timing of these oscillations was consistent with a feedforward flow of information from IT to PFC (two-way ANOVA, *p* < 0.05 for main effect of direction, *p* > 0.05 for animal). Similarly, the receipt of the reward (right column) was accompanied by a strong increase in alpha-band (8–12 Hz) synchronization that was localized to the trials near *N_50_*. This synchronization was more consistent with feedback transmission (ANOVA, *p* < 0.05 for direction, *p* > 0.05 for animal). These two synchronization events were the strongest candidates for a neural correlate of the “moment of insight” in our data.

Indeed, across sessions, synchronization between IT and PFC accurately predicted changes in behavioral performance. As shown in Figure 3B-C, there was a strong correlation between the trials at which synchronization peaked and the behavioral *N_50_* trials (r^2^ = 0.62 for gamma synchronization in the Foraging epoch and r^2^ = 0.73 for alpha synchronization in the Reward epoch, Pearson correlation tests, *p* < 0.05, corrected). These associations were significant for each animal individually (linear regression, trial ~*N_50_*, *p* < 0.05, corrected) and when the data were combined across animals (linear regression, trial ~*N_50_*, *p* < 0.05, corrected). In contrast, the absolute magnitude of the change in synchronization did not predict the behavioral moment of learning for either frequency band or epoch (two-way ANOVAs, *p* > 0.05).

Moreover, the trials with peak alpha synchronization were correlated with the trials with peak gamma synchronization (r^2^ = 0.72, *p* = 0.008), indicating that both signals were present for individual images within the same sessions (Figure S1C). Notably, we were unable to detect a timing difference between trial of peak gamma synchronization in the Foraging epoch and the trial of peak alpha synchronization in the Reward epoch (Wilks test, *p* > 0.05).

In contrast, for images that the animals failed to learn (see Methods for criteria), there was little change in synchronization across trials (two-way ANOVAs, *p* < 0.05). Thus, abrupt learning was associated with an increase in gamma synchronization in the Foraging epoch and an increase in alpha synchronization in the Reward epoch, both of which appeared to be highly specific to “moment of insight”. We therefore focus on these two signals in the following sections.

### Selection of sensory signals during abrupt visual learning

To perform the foraging task, the animals had to recognize which of the two images was present on each trial, before responding with the appropriate eye movements. According to the hypothesis outlined in the Introduction, learning to recognize images should correspond with oscillatory synchronization that is specific to sites that encode the relevant stimuli. These sites are most likely found in IT subpopulations that are selective for the images shown in each session.

To test this hypothesis, we first trained a decoder to discriminate between the two images presented in each session, using linear discriminant analysis (LDA; see Methods). The decoder was trained on multi-unit activity in IT, recorded during the Scene Onset epoch. This approach provides a close correlate of perceptual discriminability in other contexts^31^. Consistent with the importance of visual recognition for the task, more difficult image discriminations required more trials to learn: Decoding performance was negatively correlated with the number of trials required to reach *N_50_* (r = - 0.48, Pearson correlation test, p = 0.004).

To probe the cortical dynamics of this learning, we used the weights recovered from the LDA to rank each IT electrode according to the selectivity of its multi-unit activity for the two images in each session. These weights provided a measure of informativeness: Sites that were assigned high weights in the LDA were necessary for the population to discriminate between the two images, while those with low weights were not. From this analysis, we ranked IT electrodes from most informative to least informative in each session.

Figure 4 shows the average strength of interareal synchronization relative to the *N_50_* trial, computed in a moving window of five trials (see Methods) and averaged across images; data from the example image in Figure 1 are shown in Figure S1 (panel D). For visualization purposes we first focus on the 10 most and 10 least selective IT electrodes, but as shown below (Figure 6), the results are not specific to this choice.

**Figure 4.**
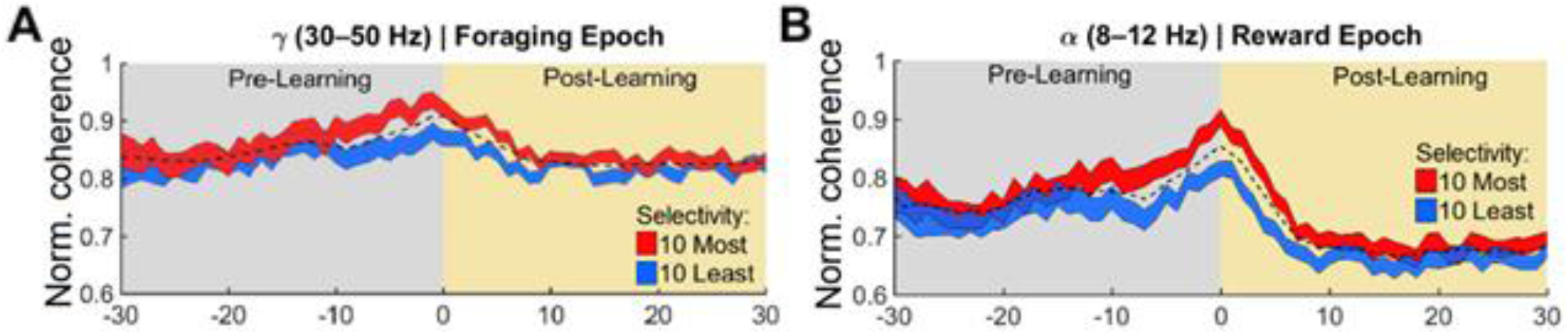
Image selective sites in IT drive synchronization around the time of learning. (A) Synchronization between IT and PFC is shown for the gamma band (30-50 Hz) in the Foraging epoch. Here data have been averaged across time within an epoch and across images. Results are aligned on the *N_50_* trial for each image. The dashed black line represents the grand median of all usable electrode pairs for all sessions. The red and blue lines correspond to the average strength of synchronization between all PFC sites and those IT sites that are most (red) and least (blue) informative about the images shown in each session. The most informative IT electrodes showed higher gamma synchronization (30-50 Hz) with PFC around the moment of learning in the Foraging epoch. Shading around each line indicates standard error (SEM). (B) As in (A), but for alpha synchronization (8-12 Hz) during the Reward epoch.

For gamma oscillations in the Foraging epoch (Figure 4A, Figure S3G), the synchronization between PFC and IT was initially similar, regardless of the informativeness of the IT electrodes. That is, early in the session, synchronization strength was similar for the most informative (red) and least informative (blue) IT electrodes. However, for later trials the most informative IT electrodes became significantly more synchronized with PFC (Figure S3G), compared to the least informative IT electrodes. A two-way ANOVA revealed a significant effect of informativeness, (*p* < 0.05) but no difference between animals and no interaction between these factors (*p* > 0.05, corrected). Interestingly, this synchronization peaked near the *N_50_* trial (Figure 4A), after which, synchronization between IT and PFC decreased gradually, with the preference for informative IT electrodes also becoming weaker. Similar results were obtained for the alpha band (Figure S3E; two-way ANOVA, *p* > 0.05) and the gamma band (Figure S3F, two-way ANOVA, *p* < 0.05) when we instead analyzed the LFP before the final saccade of each trial.

A similar pattern was seen for alpha synchronization during the Reward epoch (Figure 4B, Figure S3G). Synchronization with PFC was initially similar across IT electrodes, but as trials progressed, the strength of synchronization increased for the most informative IT electrodes, both in absolute terms and relative to the least informative electrodes (two-way ANOVA, by informativeness, *p* < 0.05, for animal or informativeness x animal interaction, *p* > 0.05, corrected). Here a distinct peak in synchronization was also evident around the *N_50_* trial, with a strong preference for informative IT electrodes. This preference also diminished after learning was complete, but it did not disappear entirely, as the increased contribution of informative IT sites to alpha synchronization persisted throughout the post-learning period (two-way ANOVA, *p* < 0.05, for animal or informativeness x animal interaction, *p* > 0.05, corrected). Similar results were evident when data for each animal were analyzed independently (Figure S2E-F). Thus, both gamma and alpha synchronization peaked around the *N_50_* trial, and long-range synchronization levels were significantly stronger for image-selective IT sites. This suggests that learning was associated with a selective flow of sensory information in both the feedforward and feedback directions.

From Figure 4A (and Figure 4B), it is also apparent that the selective contribution of different groups of IT electrodes to interareal synchronization began on average well before the *N_50_* trial. Statistically, the increased gamma synchronization of the most informative IT electrodes compared to the least informative IT electrodes began 10 trials before the *N_50_* trial (Figure S3H). For alpha oscillations, the difference emerged 14 trials before the *N_50_* trial (Figure S3G). Thus, even considering that our analysis had a resolution of five trials (see Methods), it appears that the selective synchronization of sensory signals began on average well before the behavioral moment of learning. Presently, because our analysis averages across sessions and trials, we cannot precisely quantify how abruptly or gradually this synchronization occurs.

These findings indicate that abrupt learning at the behavioral level might be caused by a synchronization of image-selective IT sites with PFC. The alternative possibility, that synchronization with PFC causes image selectivity to develop in IT, was not consistent with our data, as synchronization levels measured before learning did not predict decoding accuracy or electrode informativeness in IT, for either epoch (two-way ANOVAs, *p* > 0.05). Thus, although a conclusive test will require causal perturbations^32^, the data favor an influence of stimulus selectivity on synchronization.

This arrangement could arise trivially if unresponsive IT electrodes were included in the analysis. Unresponsive electrodes would be neither synchronized with PFC nor selective for images, and so they could exaggerate or even cause the association between synchronization and selectivity shown in Figure 4. To control for this possibility, we recomputed the synchronization analysis shown in Figure 4, after sorting IT electrodes based on the *responsiveness* of their MUA signals (Figure S2C-D, Figure S3A-B), rather than their selectivity. Responsiveness was defined as the change in MUA firing rate in the 400 ms following the release of the reward.

In contrast to the results shown in Figure 4, there was no difference in synchronization between strongly responsive and weakly responsive IT sites (two-way ANOVA, *p* > 0.05), indicating that it is image selectivity, rather than signal quality, that drives synchronization changes during learning. Consistent with this idea, the most (and least) informative electrodes changed from session to session (Figure S2A), which would not occur if defective electrodes were included in the analysis. Moreover, image selectivity by itself did not account for synchronization in other trial epochs or frequency bands (Figure S3, panels C-D).

We conclude that abrupt learning was associated with a gradual increase in synchronization between PFC and the IT sites that carried the most information about the visual stimuli. These changes were specifically related to IT selectivity, as no similar pattern was found when electrodes were sorted according to image selectivity or responsiveness in PFC (Figure S4). Thus, although previous work has found evidence of visual selectivity in PFC^33^, this selectivity does not appear to be relevant for abrupt learning in our task.

### Contribution of reward signals to abrupt visual learning

While informative sensory signals are necessary for our task, they are not sufficient, as the image identity must be linked with its corresponding RZ. The hypothesis outlined in the Introduction therefore suggests that abrupt learning might require informative sensory signals to become synchronized with neural sites that carry information about the reward context for each image.

To test this possibility, we first asked whether the location of the RZ was encoded in the neural responses in PFC. Previous work has shown that some PFC neurons respond selectively to rewarded stimuli when they are placed in specific spatial locations^34^, and so we recorded MUA responses during interleaved trials in which animals had to make a visually-guided saccade to receive a reward (see Methods). For these trials, there was no learning and no visual stimulation other than the saccade target. We then sorted the PFC electrodes according to their preference for the location of the RZ over opposite spatial locations (Figure S7I; see *Methods).*

Sites that responded selectively to the position of the RZ were not generally more coherent with IT electrodes, in either epoch or either frequency band (Figure S7). This was the case whether we defined the RZ position in retinal coordinates (relative to the fixation point on each trial) (two-way ANOVA, either retinal coordinates or by animal or the interaction, *p* > 0.05) or in screen coordinates (two-way ANOVA, either screen coordinates or by animal or the interaction, *p* > 0.05). A similar lack of RZ encoding was found in IT (Figure S7; two-way ANOVAs, *p* > 0.05). Thus, it does not appear that the sites from which we recorded play a role in maintaining the location of the RZ.

An alternative possibility is that learning in our task relies on a non-specific reward signal, as suggested by some computational models^*35*^. Indeed, a population of neurons in PFC encodes reward, independently of its associated target location^34^, and these neurons would seem to be well-suited to carry out this role. We therefore asked whether PFC sites that are sensitive to reward^36^ play a role in abrupt learning.

Figure 5 shows the LFP synchronization between PFC and IT, with data again averaged across images relative to the *N_50_* trial, as in Figure 4. However, in this case the PFC electrodes have been sorted according to the *responsiveness* of their MUAs to the release of the reward (see Methods). For alpha oscillations during the Reward epoch, the most reward-sensitive (red) PFC sites exhibited greater synchronization with IT than the least reward-sensitive PFC sites (blue) (two-way ANOVA, by responsiveness, *p* < 0.05, for animal or responsiveness x animal interaction, *p* > 0.05, corrected). In contrast to the results shown in Figure 4, this increased synchronization was not aligned on the *N_50_* trial, but rather appeared to be a non-specific signal that persisted across the duration of each session. Furthermore, in contrast to the IT results, the electrodes that provided this reward signal were consistent from image to image (Figure S3B), as would be expected based on the anatomical organization of reward signals in PFC^36^. This effect was again not due to poor electrode quality, as there was no similar preferential synchronization for other frequencies and epochs (Figure S4).

**Figure 5.**
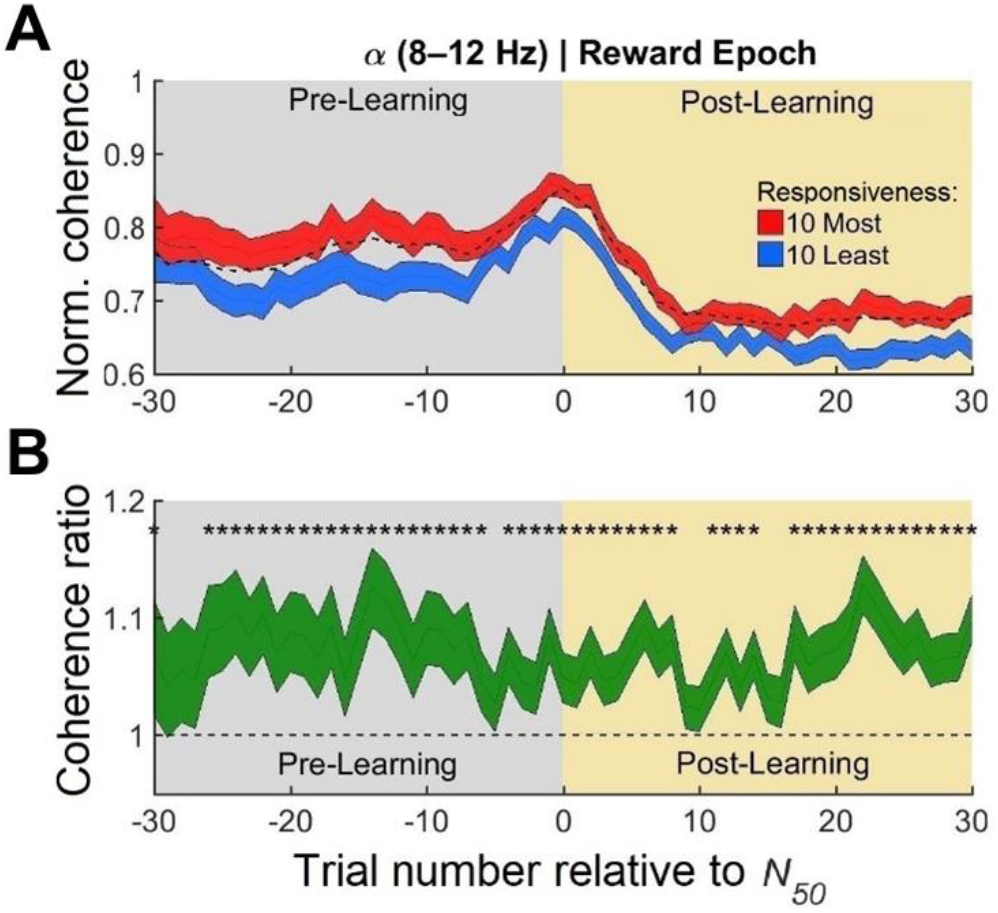
Reward responsive sites in PFC are more synchronized with sites in IT. (A) Synchronization between IT and PFC is shown for the alpha band (8-12 Hz) in the Reward epoch. As in Figure 4, data have been averaged across time and across sessions, and results are aligned on the *N_50_* trial for each image. The dashed black line represents the grand median of all usable electrode pairs for all sessions. The red and blue lines correspond to the average strength of synchronization between all IT sites and those PFC sites that respond most (red) and least (blue) strongly to the receipt of reward. The most reward-responsive PFC electrodes showed higher alpha synchronization with IT, across all trials relative to the onset of learning. Shading around each line indicates standard error (SEM). (B) The ratio of IT-PFC synchronization strength between reward-responsive and unresponsive PFC electrodes. Asterisks indicate a ratio significantly different from 1.

The full interaction between reward and sensory signals is shown in Figure 6, for all sites in both areas. Here each panel corresponds to the data averaged across images for trials relative to the *N_50_* trial. As in Figures 4 and 5, the IT electrodes have been ranked according to their informativeness about the images in each session, and this ranking is indicated on the x-axis of each panel. The PFC electrodes have been ranked according to their responsiveness to rewards, and this ranking is indicated on the y-axis of each panel. The color at each point in each map indicates the average strength of synchronization between electrodes of the corresponding ranks. Reddish colors indicate strong synchronization, and bluish colors indicate weak synchronization.

**Figure 6.**
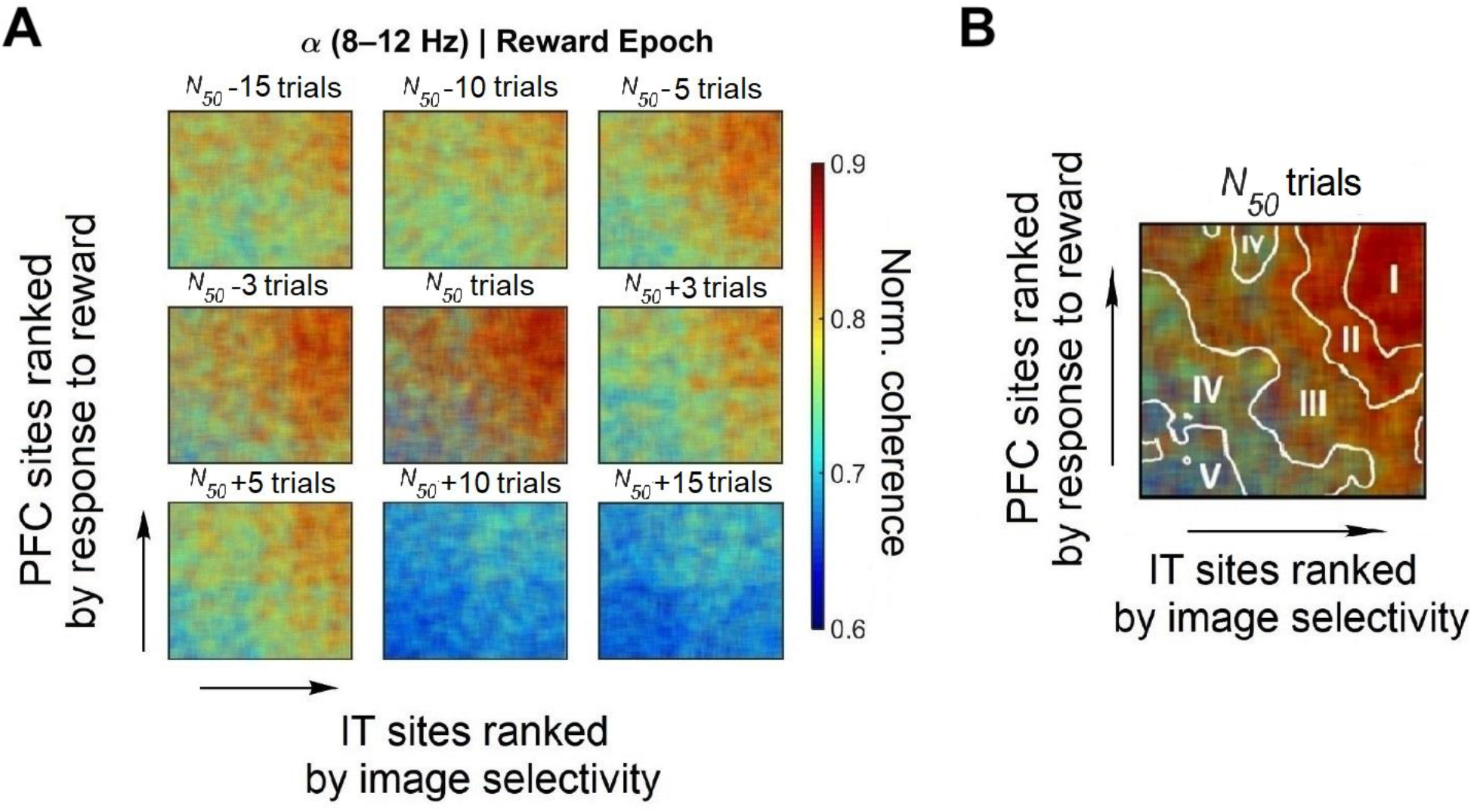
Interaction of reward responsive sites in PFC and image-selective sites in IT. (A) Alpha synchronization in the Reward epoch, for all IT and PFC electrodes across sessions. Reddish colors indicate strong synchronization, while bluish colors indicate weak synchronization. Different panels correspond to different trials relative to the *N_50_* trial. Electrodes in IT were ranked based on much information they carried about images within each session (increasing informativeness from left to right). Electrodes in PFC were ranked based on how strongly they responded following a reward (increasing sensitivity from left to right). Synchronization is generally weak and lacks organization early in each session (top left) but grows stronger around the moment of learning (central panel). Post-learning, synchronization again becomes weaker (bottom right). (B) Synchronization around *N_50_* is clearest between IT electrodes that encode images and PFC electrodes that encode rewards (top right of central panel). Data were placed into quintiles according to synchronization strength, and contours were drawn between levels to illustrate this synchronization structure.

For the trials near *N_50_* (Figure 6A, center panel) the strongest alpha synchronization is indeed limited to image-selective sites in IT and reward-sensitive sites in PFC. This is shown by the concentration of reddish colors in the upper right of the panel; the contour plot in Figure 6B emphasizes that the strongest synchronization occurs between these same subpopulations. For trials long before or after the *N_50_* trial, synchronization is generally weaker and distributed more diffusely across sites in both areas. These findings were stable across recording sessions (Figure S2J; R^2^ = 0.05, *p* = 0.09).

### (Lack of) local correlates of abrupt learning

The results thus far indicate that synchronization between IT and PFC is a strong correlate of abrupt learning. For slower forms of learning, previous work has found changes in selectivity at the single-neuron level in both IT^37, 38^ and PFC^17^. We therefore looked for similar changes in each area that might reflect abrupt learning.

We used the decoding approach outlined above to assess whether multi-unit population activity showed improved discrimination ability with learning. Decoders were trained on the same images during the Scene Onset epoch for the pre-learning and post-learning stages of each session. The results reveal that image decoding performance was generally strong in both areas, with the mean cross-validated classification rates being 87.5% (*SE*= 2.46%) in IT, and 76.3% (*SE*= 1.72%) in PFC. However, decoding performance did not change with learning, in either IT or PFC (Figure 7A). Moreover, we did not find differences in average multi-unit firing rates between the pre- and post-learning periods in either area for either animal (Figure 7B) (*t*-tests, *p* > 0.05, corrected). In fact, the pattern of population activity in each area did not change significantly across learning epochs (two-way ANOVAs, *p* > 0.05). Thus, it does not appear that neural population activity became more selective or responsive to the images during abrupt learning.

**Figure 7.**
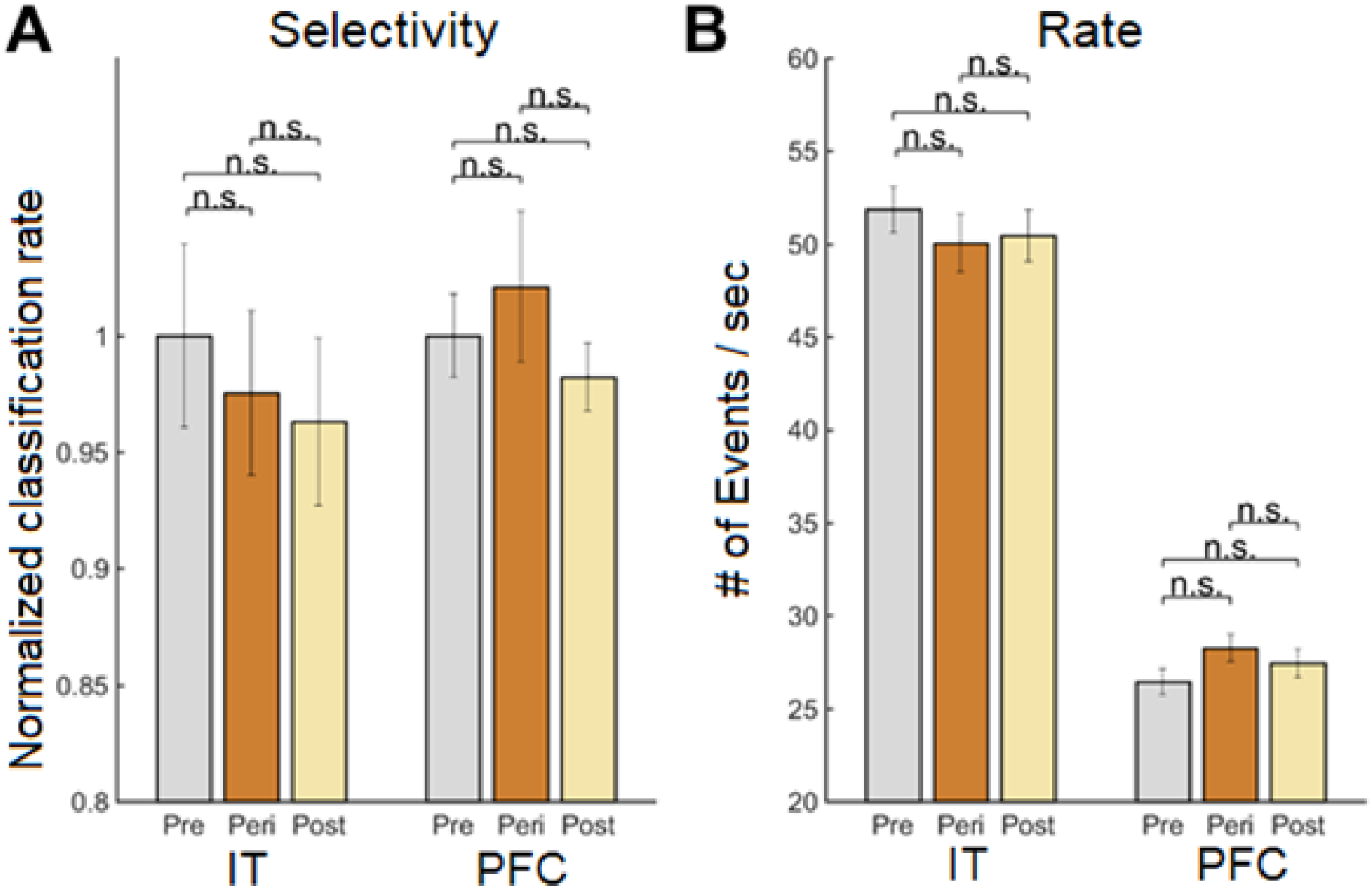
Learning does not affect multi-unit population firing activity or selectivity. (A) Selectivity of multi-units for scenes was assessed with a linear discriminant analysis (LDA), trained on the first 400 ms of spiking activity after scene onset. Neural data were split into pre-learning (gray), peri-learning (bronze) and post-learning epochs (yellow) for each image, using the sigmoid fits described in Figure 1 and the Methods. Selectivity levels in IT or PFC did not, on average, improve between learning epochs. Error bars reflect standard error (SE) across sessions. (B) Similarly, learning did not result in changes in responsiveness between learning epochs.

Similarly, there was no consistent change in LFP power (Figure S6, panels A-B) or intra-areal synchronization (Figure S6, panels C-D) around the moment of learning, either globally per area or when electrodes were sorted by informativeness in any frequency band for either animal (*t*-tests, *p* > 0.05, corrected). In IT, there was a weak negative correlation between image selectivity and both alpha (r^2^ = 0.03, Pearson correlation test, *p* < 0.01) and gamma (r^2^ = 0.04, Pearson correlation test, *p* < 0.01) LFP power (Figure S6, panels E-F). Finally, IT electrodes’ responses to reward did not change with learning in a way that was either frequency-specific, or specific to their informativeness about the image (*t*-tests, *p* > 0.05, corrected).

Across trials, there was however a steady decrease in overall PFC power and intra-areal coherence. Similar results have been reported previously, and it has been suggested that they might reflect a weaker^39^ or more selective engagement with sensory information as learning progresses. The latter possibility is consistent with the finding that overall synchronization often decreases after learning (Figure 4), while the relative advantage for image-selective sites persists (Figure S3, panels G-H). A similar conclusion has been reached from fMRI studies of learning^40^.

## DISCUSSION

Visual learning has traditionally been studied in laboratory tasks that involve extensive training to associate specific stimuli with specific responses. Such training can profoundly alter the underlying cortical circuitry^41^ but it does not necessarily reflect the kind of learning that occurs in natural settings. We have therefore examined learning in a more naturalistic, free-viewing task that approximates the kind of foraging that primates perform daily in the wild. This behavior is thought to have driven the evolutionary expansion of PFC, as well as its connection with IT^42^.

We found that non-human primates engaged in our foraging task abruptly learned to recognize visual images and to associate them with specific reward locations (Figure 1). The strongest neuronal correlates of this learning were found in the synchronization of LFPs between PFC and IT (Figure 3), and this synchronization seems well-suited to link informative visual inputs with signals about rewarded outcomes (Figures 4-5). The strength of this synchronization was greatest around the moment of insight (Figures 4-5), but the tendency for informative IT signals to be more synchronized with PFC persisted well into the post-learning phase of each session (Figure 4). Local correlates of learning were largely absent in both IT and PFC (Figure 7). These results therefore indicate that long-range synchronization is capable of supporting abrupt visual learning.

### Theories of abrupt learning

Hebb was among the first to evince a theory of abrupt learning, hypothesizing that the underlying neurophysiological mechanisms share some properties with those that support learning on longer timescales^9^. More recent work has provided some support for this idea, by showing that rapid learning exhibits stimulus specificity similar to that observed with slower learning^6, 43^.

Computational work has explained both types of learning based on a process in which decision-making structures search for the most informative sensory neurons for a given task^44, 45^. This proposition is entirely consistent with the current results (Figure 4), as well as previous findings in other brain regions^41, 46^. Moreover, in accordance with psychophysical results, we find that the selection of informative sensory inputs relies on explicit feedback^43^ in the form of rewards.

For slower forms of learning, there has been some question about whether reward signals reaching sensory cortex are stimulus-specific^47^. Our data show that the synchronization for reward-related signals is non-specific with respect to the location of the RZ (Figure S7) and unchanged by learning, being roughly constant throughout the duration of each session (Figure 5). However, the moment of insight is accompanied by a marked increase in sensory specificity: The greatest synchronization occurs for image-selective sites in IT (Figure 6). One explanation is therefore that, during learning, non-specific feedback about rewards is amplified according to the informativeness of the sensory sites for the perceptual task^48^.

The mechanism by which reward signals are connected to informative sensory signals is likely related to that of top-down attention, which is important for most kinds of learning^1^. The interplay of these different kinds of feedback is captured effectively by models that typically operate on longer timescales^49^. In these models, a successful trial (i.e., one followed by reward) is followed by an attentional signal that selectively targets the upstream neurons that led to the rewarded outcome.

Our data (Figure 3) are consistent with this idea if we consider gamma synchronization between prefrontal and visual cortices to be a correlate of covert attention, as in previous work^50^. Although we have not controlled attention directly in our experiments, a speculative interpretation of our data is that learning occurs when attention is directed to the most informative visual features in each image. This would account for the segregation of IT sites based on informativeness that occurs around the moment of learning (Figure 4). The most informative signals in IT would then propagate up the hierarchy to PFC, which is consistent with the relative (feedforward) timing of signals we observed, as well as much previous work^51, 52^.

### Role of oscillations in abrupt learning

Hebb’s favored mechanism for learning took the form of cell assemblies that become connected after being repeatedly activated at the same time. Modern investigations of this hypothesis in humans have confirmed that long-range synchronization between different brain regions is a reliable correlate of certain kinds of learning^53, 54^. This coherence has often been found in the gamma band^13, 53^, which is also associated with learning in our task (Figure 3, Figure 6). Although the relationship between gamma oscillations and learning was correlational in this work, there is some evidence for a causal connection as well^21, 55^.

In rodents, gamma oscillations are often nested within theta oscillations, and this coupling has been shown to be important for associative learning^56^. Moreover, the theta rhythm itself is well-known for its role in learning, particularly in the hippocampus^57, 58^, but theta did not seem to correlate with abrupt learning in our data (Figure 3). Rather, the clearest correlate was found for PFC-driven oscillations in the alpha band (Figure 4), which are also implicated in human studies of insight^55^.

We found that alpha synchronization seemed most related to the release of rewards within each trial (Figure 5). This makes sense in light of previous findings, in which alpha oscillations in PFC are modulated by dopaminergic signals arriving via the striatum^59^. Indeed, blocking dopamine receptors in PFC impairs associative learning and alters alpha oscillations, and as a result it has been suggested that alpha plays a role in controlling neural selectivity for sensory inputs^59^. More generally, alpha is commonly thought to gate neural activity according to task demands^60^. Our data suggest that these signals flow in a feedback direction from PFC to visual cortex, given their frequency distribution^51, 61^ and their timing relative to IT.

### Abrupt learning in the prefrontal cortex

Most previous work has suggested that PFC plays a role in facilitating learning but that it is not itself the primary site of long-lasting plasticity^62, 63^. Instead, it might be involved in discerning the rules that map stimuli to responses for a given task^64^. The encoding of these rules can appear abruptly in PFC for novel associations^65^ or for reversals of previously learned associations^66^. However, learned improvements in stimulus encoding are less common, and there is even evidence that stimulus encoding decreases with learning^39^, suggesting that PFC likely does not learn novel stimuli on short timescales. Our results (Figure 3, Figure 4) are consistent with this idea, since there is little evidence of enduring changes in image selectivity in the PFC neurons we recorded (Figure 7).

For visuomotor associations, the connection between PFC and IT is known to be critical^19^. Lesions that disconnect these areas severely impair visuomotor learning and especially “object in place” learning, a kind of rapid learning that resembles our foraging paradigm^19^. A more mechanistic role for PFC is suggested by the finding that neurons in this area help to solve the “credit assignment” problem. This refers to the challenge faced by the brain in identifying synaptic connections that can be strengthened to improve performance on a given task. Asaad et al. (2017) reported that selective visual signals were found in populations of PFC neurons that also represented reward outcomes, as would be necessary to solve the credit assignment problem^67^. Our results suggest that this information emerges dynamically through communication between subpopulations of IT neurons that encode the relevant stimuli and PFC neurons that encode the reward (Figure 6).

### Abrupt learning in the visual cortex

While previous work has found rapid learning in individual IT neurons^38, 68^, we did not find evidence for this at the population level (Figure 7). One possibility is that learning affected different classes of neurons differently^69^, which would not have been reflected in our population-level results, as we do not have a reliable way of assessing molecularly-defined cell types in our sample.

With extended training, changes in neural selectivity have been observed in IT^70, 71^, V4^72, 73^, V2^4^, and V1^5^. These results are generally consistent with the observation that the adult visual system changes quite slowly with training^1^. Far less is known about the effects of rapid learning in the visual cortex, but Wang and Dragoi (2015) reported a transient, learning-related increase in spike-field coherence in area V4^74^. This coherence increase was found in the theta band and within a single area but followed a similar time-course to the changes in long-range synchronization we report here (Figure 3).

### Other brain structures involved in abrupt learning

The most dramatic form of abrupt learning is one-shot learning, in which only a single instance of a stimulus is provided^75^. This kind of learning is closely related to episodic memory, which depends critically on the hippocampus^76^. Interactions between PFC and hippocampus have been shown to correlate with learning on short timescales in monkeys^28^ and rodents^57^. In humans, the hippocampus is recruited by PFC when one-shot learning is dictated by the demands of a task^77^. Thus, it might be that PFC establishes the conditions for learning, but that information is initially stored in the hippocampus^78^ and then later consolidated into long-term memory^79^.

For tasks that involve a saccadic response, neurons in the caudate nucleus facilitate saccades toward rewarded locations, and these signals can update rapidly as contingencies change^66, 80^. Indeed, Williams & Eskandar (2006) found that firing rates in the caudate correlate with the rate of learning in a task that requires learning to saccade to a particular visual stimulus^81^. The output of these neurons could therefore be linked to the synchronization changes we have observed (Figure 4), which also correlate with the rate of learning.

This connection between dopaminergic cells and learning in PFC has been studied by Puig and Miller (2012), who showed that dopaminergic drugs in PFC can modulate associative learning^59^. Specifically, antagonists of some dopamine receptors block the formation of new associations, but they do not impair the recall of those already established. Putting these results together, it could be that dopamine released from the caudate facilitates the formation of new or stronger connections between informative sites in IT and longer-term storage in hippocampus, with the PFC playing a permissive role in “switching on” rapid learning^62, 82^. Previous work has also highlighted a role for the amygdala in this process^22, 83^.

Rapid learning has also been found in auditory tasks^84^. For example, observers exposed to meaningless sounds learn to recognize them after repeated exposures; as in our foraging task, this kind of learning occurs abruptly and endures for several weeks afterwards^85^. Learning in both types of tasks might reflect a general mechanism for assimilating statistical regularities about the environment^86^.

### Alternative interpretations

In addition to providing signals about reward outcomes, the PFC plays a role in arousal, working memory, and attention^36^, all of which are important for learning^1^. Though it is notoriously difficult to disentangle these factors^87^, we have shown several lines of evidence suggesting that none of them alone are sufficient to explain our findings. First, non-specific effects like arousal would not account for the preference for informative IT electrodes in the synchronization between IT and PFC (Figure 4). Thus, while we favor the interpretation of a non-specific signal emanating from PFC (Figure 5), we suggest that it is more parsimonious to think of it as carrying information about rewards, rather than arousal *per se*^35^. Second, working memory might account for the retention of visual associations within each testing session, and it has been often associated with both gamma^88^ and alpha synchronization^89, 90^. However, by itself it cannot account for the animals’ ability to remember the associations many days later.

This long-term recall also supports the idea that changes in behavioral performance in our foraging task reflect genuine visual learning. This is in contrast to an alternative possibility, namely that the data reflect a shift in behavioral strategy between early and later trials. Such shifts might be expected to alter the dynamics of prefrontal activity^91^, but it is not clear why they would lead to the pattern of synchronization changes we have observed (Figure 3), why this pattern would emphasize informative IT electrodes (Figure 4), or why it would have effects that persisted many weeks later.

Finally, we have not specifically controlled top-down attention in this task, as part of our goal was to simulate learning under natural conditions. Attention could therefore have played various roles in learning or overall task performance^92^. Of these roles, the most commonly observed consequence of attention is an improvement in sensory coding, which we did not observe in our data (Figure 7). However, our data are consistent with a different role for attention, namely that of selecting relevant visual inputs based on rewards^92, 93^. As mentioned above, such a role is consistent with both the gamma synchronization we observe^50^ and its specificity for informative sensory signals (Figure 4). Previous work has found that this kind of attentional mechanism is especially important for extracting relevant information from complex stimuli^94, 95^ of the kind we have used in our foraging task, and so it could have been a factor in our data as well.

## CONCLUSION

We have shown that visual learning happens abruptly when animals use eye movements to forage for a rewarded location in an image. In contrast to most previous studies of visual learning, our paradigm made use of natural images and unconstrained eye movement behaviors. As a result, our results might provide greater insight into the kind of learning that occurs outside of a laboratory. Moreover, in contrast to slower forms of learning, our paradigm readily facilitates observation of neural signals throughout the entire trajectory of learning.

In our electrophysiological data, abrupt learning was accompanied by increased synchronization between reward signals in PFC and image-selective signals in IT. Time-frequency analysis suggests that this synchronization has two components: a feedforward component that is active during inspection of the image and a feedback component that is active upon receipt of a reward. This pattern of activity is compatible with predictions of computational models that assign specific functions to feedforward and feedback pathways during visual learning^49, 96^. Both signals become more attuned to informative sensory signals at the “moment of insight”, highlighting the role for distributed neural circuits in facilitating the rapid acquisition of new sensory knowledge.

## STAR★METHODS

Detailed methods are provided in the online version of this paper and include the following:

- KEY RESOURCES TABLE
- RESOURCE AVAILABILITY
  - Lead contact
  - Materials availability
  - Data and code availability
- EXPERIMENTAL MODEL AND SUBJECT DETAILS
  - Animals
- METHOD DETAILS
  - Implant preparation and surgical procedures
  - Behavioral task
  - Eye calibration
  - Saccadic eye movements
  - Processing of raw neural data
  - Trial epoch selection
  - Calculation of spectral power
  - Synchronization
  - Calculation of spectral Granger-Geweke causality
  - Multi-unit analysis
- QUANTIFICATION AND STATISTICAL ANALYSIS
  - Response modeling
  - Statistics

## SUPPLEMENTAL INFORMATION

Supplemental Information is available online at T.B.D.

## ACKNOWLEDGEMENTS

This work was funded by a grant from the CIHR to C.C.P. (PJT-461642) and a graduate student fellowship to B.A.C. from NSERC. We thank Dr. Shahab Bakhtiari and Tugce Gurbuz for helpful comments and discussions.

## AUTHOR CONTRIBUTIONS

M.R.K. and C.C.P. designed the study. M.R.K. and T.P.Z. performed the experiments. B.A.C. analyzed the data. B.A.C., M.R.K. and C.C.P. wrote the paper.

## DECLARATION OF INTERESTS

The authors declare no competing interests.

## STAR★METHODS

### KEY RESOURCES TABLE

| REAGENT or RESOURCE | SOURCE | IDENTIFIER |
| --- | --- | --- |
| **Experimental Models: Organisms/Strains** |  |  |
| Rhesus macaque, *Macaca mulatta* (males, 4 and 10 years) | McGill University | N/A |
| **Software and algorithms** |  |  |
| MATLAB R2016b | Mathworks | RRID:SCR_001622 |
| Chronux | http://chronux.org<sup>97</sup> | v2.12.3 |
| Microsaccade Detection | (Otero-Millan et al., 2014)<sup>98</sup> | N/A |
| 3D Slicer | http://slicer.org<sup>99</sup> | 4.11 |
| Custom code | This paper | https://github.com/bennettcsorba/AbruptLearning |
| **Other** |  |  |
| Utah multielectrode arrays | Blackrock Microsystems, Utah (USA) | Iridium Oxide, 96 Channel |
| Macaque Atlas (Figure 2) | (Markov et al., 2014)<sup>30</sup> | Core-Nets<br>https://core-nets.org |

### RESOURCE AVAILABILITY

#### Lead contact

Requests for information or resources should be directed to the lead contact, Bennett Csorba.

#### Materials availability

This study did not generate any unique materials.

#### Data and code availability

- All behavioral and neural data used in this study are available from the lead contact upon reasonable request (due to the size of the data set).
- All custom MATLAB scripts related to this paper are available on GitHub (https://github.com/bennettcsorba/AbruptLearning) and are publicly available as of the date of publication.
- Any additional information required to reanalyze the data reported in this paper is available from the lead contact upon request.

### EXPERIMENTAL MODEL AND SUBJECT DETAILS

#### Animals

Two adult male rhesus macaque monkeys (*Macaca mulatta,* 4 and 10 years old) participated in the study. All animal care, surgical and experimental procedures were approved by the Montreal Neurological Institute’s Animal Care Committee, and the animals were monitored by the veterinary staff for the duration of the experiment.

### METHOD DETAILS

In some experiments on the same animals, we examined the influence of transcranial direct current stimulation (tDCS) on neural activity and behavior. These data have been reported in a separate paper^21^ and were excluded from the analyses reported here.

#### Implant preparation and surgical procedures

Experimental preparation has been described in a previous publication^21^. In brief, high-resolution T1 and T2-weighted anatomical MRIs (0.6-0.8 mm^3^ voxels) were acquired for each animal and were used for surgical planning. Animals were implanted with a custom titanium head post (Hybex Innovations, Montreal Canada) using standard sterile surgical techniques. Post-recovery, animals were familiarized with the laboratory environment, and were trained for head fixation and subsequently the behavioral “foraging” task. Once overall performance in this paradigm was stable, we prepared the animals for electrophysiology.

A second surgery was then performed to implant multielectrode arrays (Blackrock Microsystems; Utah, USA) into IT (area TEO) and the lateral PFC, ventral to the principal sulcus (area 46v). This part of PFC receives projections from IT and is often considered part of the machinery for visual image recognition. A neuro-navigation system (Brainsight Vet; Rogue Research Montreal Canada) synced with previously completed MRI scans^99^ was used to ensure that the arrays were accurately positioned. Arrays were then inserted into PFC and IT using a pneumatic device. The positions of the arrays were verified post-operatively (Figure 2), via coregistration of preoperative MRI scans with postoperative CT scans.

#### Behavioral task

Non-human primates were seated in front of a screen that spanned 30×60 degrees of visual angle, at a viewing distance of 40 cm. Eye positions were sampled at 500 Hz with an infrared eye tracker (SR Research, Ontario). During each session, the animals completed the foraging task described below over the course of approximately 30 minutes, with each trial following the progression shown in Figure 1A. Foraging trials were alternated with brief trials involving a calibration task (see below), to ensure precise calibration of the eye tracker. The foraging paradigm was adapted from a previous study which investigated associative learning in humans^20^.

Each trial of the foraging task began with the appearance of a high-contrast fixation cross on a gray screen. Animals initiated the trial by fixating within 2 degrees of the fixation spot for 750–1000 ms, after which they were presented with a full-screen image. Images were chosen from collections of Creative-Commons and public domain photographs of natural images (url: flickr.com). Image presentation order was randomized within each session, and sessions consisted of 75 or 100 trials (depending on the animal) of each of the two images.

A 2° reward zone (RZ) was embedded into a random location in each image, and jittered a small amount on each trial, according to a bivariate normal distribution with a standard deviation of 1–4 degrees. This encouraged the subjects to learn the spatiotopic location of the RZ, rather than a specific set of saccade vectors (Figure S1, panels G-J), as the RZ was positioned differently relative to the starting eye position on each trial (Figure S1, panels K-L). Upon image presentation, animals could freely search the image until they found the RZ or 15 (Monkey F) or 20 (Monkey M) seconds elapsed. If animals did not successfully find the RZ within that time, a high-contrast cue was shown at the RZ to guide their saccades into it. Once the animals had successfully maintained gaze position within the RZ for 100 ms, they received a juice reward and the trial ended.

Throughout our electrophysiological recordings, behavioral performance was not associated with session number (linear regression between session number and *N_50_*, R^2^ = 0.04, p = 0.12), as animals were trained extensively on the task prior to the commencement of neural recordings. Thus, any fluctuations in performance across sessions are likely due to differences in image difficulty or engagement.

In the main experiment, images were not repeated across sessions. We also conducted 19 long-term recall sessions, in which animals “foraged” in image-reward zone pairs that they had seen 3–128 days earlier. Behavioral performance during these sessions is described separately in the Results, but the data are not included in any of the electrophysiological analyses.

#### Eye calibration

Eye calibration was checked on interleaved trials in which the animals were shown a grey screen with a single small, high-contrast saccade target, randomly located at one of 9 (or 25) locations on a 3×3 (or 5×5) grid spanning the central 14 horizontal and vertical degrees on the monitor (Figure S7I). Animals received a liquid reward for making a saccade to the target and maintaining their gaze on it for 750–1,250 ms.

#### Saccadic eye movements

Saccade onset and offset were estimated from the eye position traces based on eye velocity and acceleration^100^, followed by manual review to make small corrections to onset/offset times and to discard false detections. Microsaccades were extracted using an unsupervised clustering method^98^.

#### Processing of raw neural data

Wideband neural signals were recorded using a neural interface processor (Ripple Neuro, Salt Lake City, Utah). Neural signals were sampled at 30,000 Hz and band-pass filtered between 0.3 and 7,500 Hz during acquisition. These data were post-processed offline to remove powerline and low-frequency movement artifacts, using the Chronux^97^ functions `rmlinesc` and `locdetrend` respectively. Next, the local field potentials (LFPs) were extracted with a fourth order Butterworth low-pass filter (F_c_ = 500 Hz) and re-sampled at 1 kHz.

Subsequent analysis was performed with the MATLAB signal processing package Chronux (v2.12.3) and custom-written MATLAB software. The LFPs were manually reviewed for quality. Sites were excluded from analysis on individual sessions when they had an exceptionally low signal-to-noise ratio (SNR < 3). An average of 89 out of 96 IT sites, (*SE*=0.33, > 92% of all sites) and 95 of 96 PFC sites (*SE*=0.12, > 98% of all sites) were available for analysis.

#### Trial epoch selection

Three epochs of interest (Figure 1A) were chosen from the data from each trial: a Scene Onset epoch (400ms following scene onset), a Reward epoch (400ms following reward onset), and a Foraging epoch (225ms to 100ms before the onset of the first saccade on each trial). We chose to limit the analysis of the Foraging epoch to the time just before the onset of the first saccade because the free-viewing paradigm rendered the Foraging epoch otherwise uncontrolled in terms of eye movements and visual stimulation. In particular, eye movements can strongly influence LFP signals^58, 101^.

The epoch durations were chosen based on preliminary analyses (Figure 3), with the intention to identify regions of interest in time-frequency space. Other bands and window sizes around these event epochs were considered for all reported neurophysiological measures to avoid biasing the results. See Figure 3, Figure S3 and Figure S4 for other frequency bands, which showed no effect.

#### Calculation of spectral power

Within each epoch, we estimated the LFP power with multi-taper methods, using five tapers and a spectral resolution of 7.5 Hz^102^ (Figure S6, panels A-B).

For both recording areas, we took the average power across all usable electrodes on each trial, within each trial epoch. For each trial epoch, a maximal trial value was determined, to which all other trial values were normalized. Learning epoch averages were then computed from the normalized values.

#### Synchronization

To quantify synchronization (Figure 3–6, Figure S1 panels C-D, Figures S2 panels C-H, Figure S3, Figure S4, Figure S6 panels C-D, Figure S7 panels A-H), we computed LFP-LFP coherence, which is a measure of the linear correlation between two signals in the frequency domain^103, 104^ and a commonly-used metric of synchronization^104^. Coherence is independent of absolute phase and is often thought of as the consistency across trials of the relative phase angles between two signals. We computed magnitude-squared LFP-LFP coherence with multi-taper methods, using the Chronux^97^ functions cohgramc and coherency for continuous signals. Five tapers and a spectral resolution of 7.5 Hz^102^ were used. The magnitude-squared coherence between two signals at a given frequency is defined as the cross-power spectrum (also called cross-spectral density) of those two signals divided by their individual power spectra, also taken at that same frequency.

Session averages for LFP-LFP coherence (Figure 4) were computed by taking the median values across usable electrodes within (max. 4,560 pairs each) or between (max. 9,126 pairs) IT and PFC. Because coherence is not defined on a single trial, we used five-trial windows to compute coherence across trials throughout each session. This allowed us to capture relatively abrupt changes in the communication between IT and PFC that occurred over the course of a few trials. The first and last two trials of each session used three-trial and four-trial windows respectively to compute coherence, but these data were rarely used.

As with power, we normalized coherence values to the maximum values within each trial epoch. Normalization was done independently at each frequency. Averages within each learning epoch (Figure 3, Figure S1C) were then computed from the normalized trial values for each trial epoch.

Correlations between coherence levels and behavior (Figure 3B) were obtained by subsampling different groups of electrodes in each area 1000 times and computing the linear regression between the time of peak coherence and the *N*_50_ trial for the corresponding session.

Mean values of r^2^ ranged from 0.39, for 10 electrodes per area, to 0.73 for 96 electrodes per area. Only the latter value is reported in Figure 3B.

#### Calculation of spectral Granger-Geweke causality

Within individual sessions, non-parametric spectral Grange-Geweke causality estimates^105^ were computed between all usable electrode pairings, using trial data from three learning epochs for frequency bands of interest. Causal estimates were then averaged by learning epoch across all IT-PFC electrode pairings. The size of the temporal window used in each trial was dependent on the trial epoch (see above).

#### Multi-unit analysis

Because single-unit activity was generally sparse in both areas, we focused on multi-unit activity in this study (Figure S5). A three-step process was used to extract multi-unit activity from the wideband signals. First, the raw signals were high-pass filtered (F_c_ = 500 Hz), using a 3rd-order Butterworth filter. Multi-unit events were then defined as the instants where the resulting signal exceeded 3 robustly-estimated standard deviations from the mean (i.e., 4.2 median absolute deviations). To avoid double-counting events, we imposed a 2-ms refractory period immediately after each threshold crossing. We defined MUA responsiveness as the mean rate of MUA events (Figure 7) in units of events/s, occurring on each electrode during the 400 ms after the reward was dispensed (“reward responsiveness”) or after scene onset (“image responsiveness”).

We used a linear discriminant analysis (LDA) to identify subsets of electrodes whose multi-unit activity were more (or less) selective for the scenes being presented, and to determine if neuronal selectivity improved with learning. Classification was done independently for each experimental session, and classifiers were trained separately on IT activity and PFC activity using scene labels as class inputs (Figure 7). Two unique scenes were shown on every session, so the classifier was always trained to discriminate between two data classes. LDA classifiers were trained on multi-unit activity collected on all available electrodes in IT and PFC, using the responses obtained at each site during the scene onset epoch. We chose this epoch because it provided the purest measure of sensory selectivity, without confounding influences of eye movements or rewards.

Classification performance on individual sessions was computed using leave-one-out cross-validation. Because population selectivity did not change post-learning in either IT or PFC (Figure 7A), the classifiers were trained on all trials within a session. Classifiers were also trained on shuffled class data as a control, where the performance was approximately 50%, as expected for a balanced two-way classification task. Classifiers consistently relied on different IT electrodes to classify images across sessions (Figure S2A) in each animal, as would be expected from the variable selectivity of different IT sites^32^.

To determine whether MUAs encoded the location of the RZ, we first identified saccades in the eye calibration trials made to targets located at similar spatial coordinates to the RZ (Figure S7I) and its reflection across both axes. This was done by matching the RZ coordinates for each image to the closest target location (by Euclidean distance) in the calibration task. The median Euclidean distance between the RZ and the closest target location in the calibration task was 2.97 degrees across images. One image whose RZ had both an absolute azimuth and elevation of less than 2 degrees was excluded from this analysis, since the RZ’s spatial reflection was directly adjacent to it.

We then used activity recorded during the calibration trials to estimate the response of each MUA signal, during the 250 ms before the onset of the saccade to the corresponding target. From these responses we calculated the preference for the direction of the RZ as a function of the multi-unit firing rates μ:

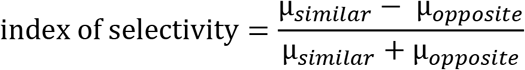

This yielded an index of selectivity, which we used to sort electrodes from most to least selective; we then analyzed interareal coherence levels for different levels of RZ selectivity in each frequency band and epoch (Figure S7). For the retinotopic analysis, the procedure was identical, except that RZs were sorted according to their position relative to the fixation point on each trial. Because the fixation point location varied somewhat from trial to trial (see Foraging task above), the retinotopic and screen coordinate systems were not identical.

To determine whether population activity changed with learning, we generated a vector of population responses for each image, averaging responses across 5 trials. We then correlated the vector with a similar vector calculated from the subsequent 5 trials and repeated this procedure throughout the course of each session.

### QUANTIFICATION AND STATISTICAL ANALYSIS

#### Response modeling

In each session, response times (RTs) for all presentations of each image were extracted. RTs were then smoothed with a three-element median filter to exclude transient lapses in engagement. The resulting response time curves were well-described by sigmoid functions, so the data were fit to a model described by the equation:

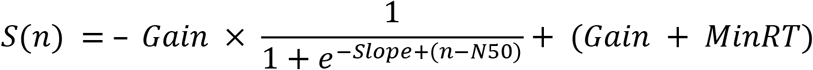

where *S*(*n*) is the response time on the *n*th presentation of a given image. The *Gain* parameter represents the magnitude of the performance improvement obtained through learning. The parameter *MinRT* reflects the animal’s asymptotic performance, or the response time necessary to complete the task once the association had been completely learned.

Similarly, the *N_50_* parameter indexes the trial number at which the response time reaches the halfway point between its initial value and its asymptotic value. This parameter therefore identifies the midpoint of the shift from a pre-learning phase to a post-learning phase (Figure 1B). The *Slope* reflects how quickly the response time shifted from its maximum to its minimum.

We excluded data from images in which the parameters of the sigmoid fits were inconsistent with reliable task performance. For example, images for which *N_50_* < 0 were generally too noisy to be interpreted. Images for which the animal failed to learn the association were those in which *N_50_* was greater than 100, *Gain* was less than 3 s, or *MinRT* was greater than 3 s. We verified that the excluded data reflected a lack of learning, in terms of the overall improvement in performance throughout the session, but we did not perform any selection based on the abruptness of learning.

#### Statistics

To ensure that the results were similar between the two monkeys, we generally used permutative two-way ANOVAs^106^, with monkey identity as a factor. Where appropriate, we used individual *t*-tests. When testing linear correlations between two groups, Pearson’s *r* values were computed.

All significance thresholds were corrected for multiple comparisons where necessary. Correcting for multiple comparisons was done by adjusting with a Holm-Bonferroni correction or a false discovery rate criterion^107^, depending on the correct context. All statistical tests were performed in MATLAB.

## Notes

### Competing Interest Statement

The authors have declared no competing interest.

### Summary of Updates

Additional control analyses added; cosmetic changes to main and supplemental figures.

