## Supplemental Materials for "Long-range cortical synchronization supports abrupt visual learning"

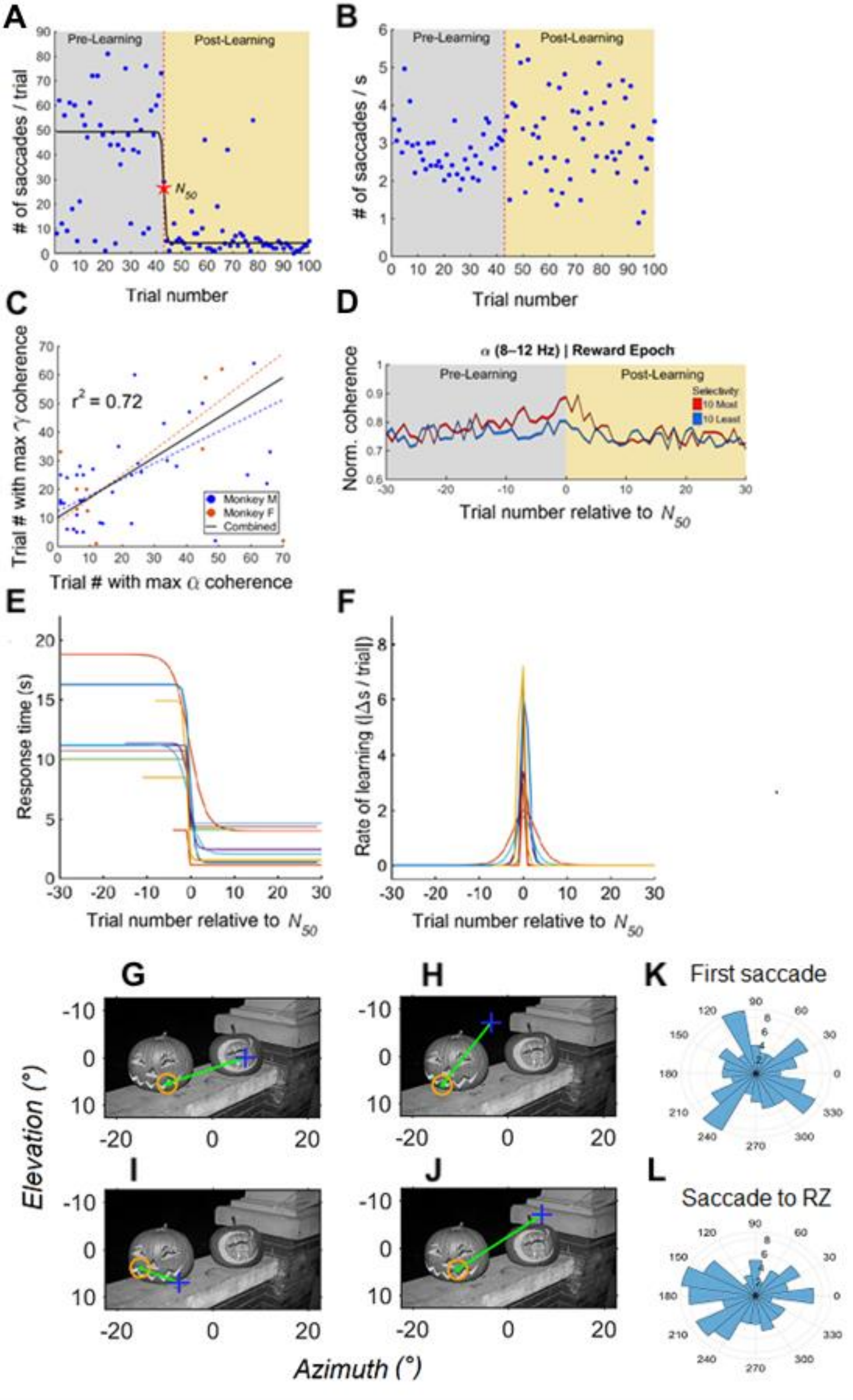
A
Pre-Learning
Post-Learning
### of saccades / trial
$N_{50}$
Trial number
B
### of saccades / s
C
Trial # with max γ coherence
$r^2 = 0.72$
Monkey M
Monkey F
Combined
Trial # with max α coherence
D
α (8–12 Hz) | Reward Epoch
Norm. coherence
Selectivity
10 Most
10 Least
Trial number relative to $N_{50}$
E
Response time (s)
F
Rate of learning (|Δs / trial|)
G
H
I
J
Elevation (°)
Azimuth (°)
K
First saccade
L
Saccade to RZ

**Figure S1, related to Figure 1 and Figure 4. Additional behavioral and neural data for the example session in Figure 1 and split for individual animals.**

(A) The number of saccades made on each trial for the example image from Figure 1 (panel B). The animal transitioned from making many saccades to few saccades to the target around the moment of learning.

(B) The rate at which saccades were made on each trial for the example image in Figure 1 (panel; B).

(C) Correlation between trial number with maximum alpha and trial number with maximum gamma coherences.

(D) Synchronization between IT and PFC is shown for the alpha band (8-12 Hz) in the Reward epoch for the example session, with data aligned on the $N_{50}$ trial. The dashed black line represents the grand median of all usable electrode pairs within the session. The red and blue lines correspond to the average strength of synchronization between all PFC sites and those IT sites that are most (red) and least (blue) informative about the images shown in each session. The most informative IT electrodes showed higher alpha synchronization (8-12 Hz) with PFC around the moment of learning in the Reward epoch, as in Figure 4 (panel B). Shading around each line indicates standard error (SEM) across electrodes.

(E) Estimated sigmoid fits for all images for Monkey F, aligned on the $N_{50}$ *value for each image*. Learning is abrupt for most images.

(F) The rate of learning, defined as the performance improvement as a function of trial number, for the images shown in E. Rates are aligned on the $N_{50}$ *value for each image.*

(G-H) Example positions of the starting eye position (blue cross) relative to the RZ (orange circle) for a single image. Trials *before* the image's $N_{50}$ trial.

(I-J) Example positions of the starting eye position (blue cross) relative to the RZ (orange circle) for a single image. Trials *after* the image's $N_{50}$ trial.

(K) For the example image from (G-J), the distribution of directions of the first saccade made on every trial.

(L) For the example image from (G-J), the distribution of directions of the final saccade made (i.e., the saccade to the RZ) on every trial.

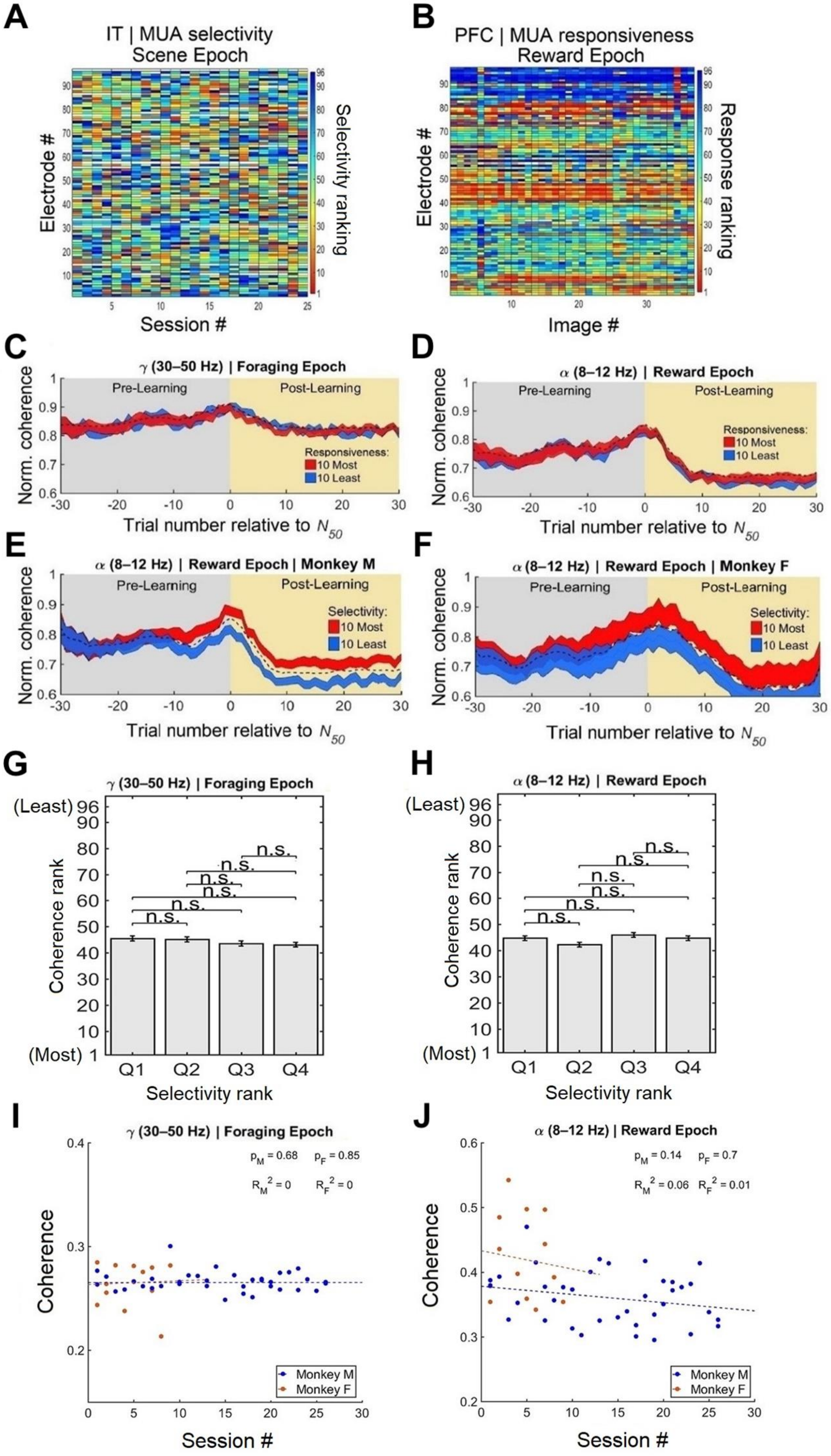
A
IT | MUA selectivity
Scene Epoch
Electrode #
Selectivity ranking
Session #
B
PFC | MUA responsiveness
Reward Epoch
Electrode #
Response ranking
Image #
C
γ (30–50 Hz) | Foraging Epoch
Pre-Learning
Post-Learning
Responsiveness:
10 Most
10 Least
Norm. coherence
Trial number relative to N50
D
α (8–12 Hz) | Reward Epoch
Pre-Learning
Post-Learning
Responsiveness:
10 Most
10 Least
Norm. coherence
Trial number relative to N50
E
α (8–12 Hz) | Reward Epoch | Monkey M
Pre-Learning
Post-Learning
Selectivity:
10 Most
10 Least
Norm. coherence
Trial number relative to N50
F
α (8–12 Hz) | Reward Epoch | Monkey F
Pre-Learning
Post-Learning
Selectivity:
10 Most
10 Least
Norm. coherence
Trial number relative to N50
G
γ (30–50 Hz) | Foraging Epoch
(Least)
(Most)
n.s.
Coherence rank
Q1 Q2 Q3 Q4
Selectivity rank
H
α (8–12 Hz) | Reward Epoch
(Least)
(Most)
n.s.
Coherence rank
Q1 Q2 Q3 Q4
Selectivity rank
I
γ (30–50 Hz) | Foraging Epoch
pM = 0.68 pF = 0.85
RM2 = 0 RF2 = 0
Coherence
Monkey M
Monkey F
Session #
J
α (8–12 Hz) | Reward Epoch
pM = 0.14 pF = 0.7
RM2 = 0.06 RF2 = 0.01
Coherence
Monkey M
Monkey F
Session #

**Figure S2, related to Figures 4, 5 and 6. Contribution of different electrodes to image selectivity and reward responses across sessions. IT responsiveness does not drive synchronization. Results are consistent across animals.**

(A) LDA selectivity rankings for every IT site are plotted for each individual session for Monkey M. Red indicates highly informative sites, while blue indicates uninformative sites. The informativeness of each site varies across sessions, indicating that different electrodes were selective for different images.

(B) MUA responsiveness to reward for every PFC site is plotted for each individual session for Monkey M. Red colors indicate highly responsive sites, while blue colors indicate sites with weaker responses. The electrodes that show responses to the reward onset are similar from experiment to experiment.

(C-D) Normalized coherence between IT and PFC is shown low-gamma (30-50 Hz) synchronization during foraging (C) & alpha (8-12 Hz) synchronization following reward (D). Each band represents the grand median of all usable electrode pairs for all images. Error bars display standard error (SEM) across images. Average levels of synchronization, with the data split by *responsiveness* to the appearance of images in IT. For all epochs and bands, there was no statistical difference in the extent to which PFC was synchronized with more responsive (red) and less responsive (blue) sites in IT. The dashed black line indicates the overall average synchronization, which did not change noticeably across trials for these frequencies and these epochs.

(E-F) Figure 4 (panel B) with data for each animal shown separately. The changes in synchronization across time, as well as the preference for informative IT electrodes, are present for each animal individually.

(G-H) Individual sites in IT were sorted based on their average coherence levels with sites in PFC (pre-learning) and checked for differences in decoding accuracy (by quartile) on individual sessions. There was no consistent statistical relationship between coherence levels and decoding accuracy for either the Foraging epoch (using gamma coherence) or the Reward epoch (using alpha coherence) (two-way ANOVAs, $p > 0.05$). Means are of all sites across sessions (Monkey M = 3247 electrodes from 37 images, Monkey F = 1194 electrodes from 13 images), and bars represent standard error (SEM).

(G) Ranking of mean pre-learning low-gamma (30-50 Hz) coherence (Foraging epoch) versus image selectivity rank for IT sites.

(H) Ranking of mean pre-learning alpha (8-12 Hz) coherence (Reward epoch) versus image selectivity rank for IT sites.

(I) Mean alpha (8-12 Hz) non-normalized coherence (early trials) between IT & PFC during the Reward epoch versus session number.

(J) Mean low-gamma (30-50 Hz) non-normalized coherence (early trials) between IT & PFC during the Foraging epoch versus session number.

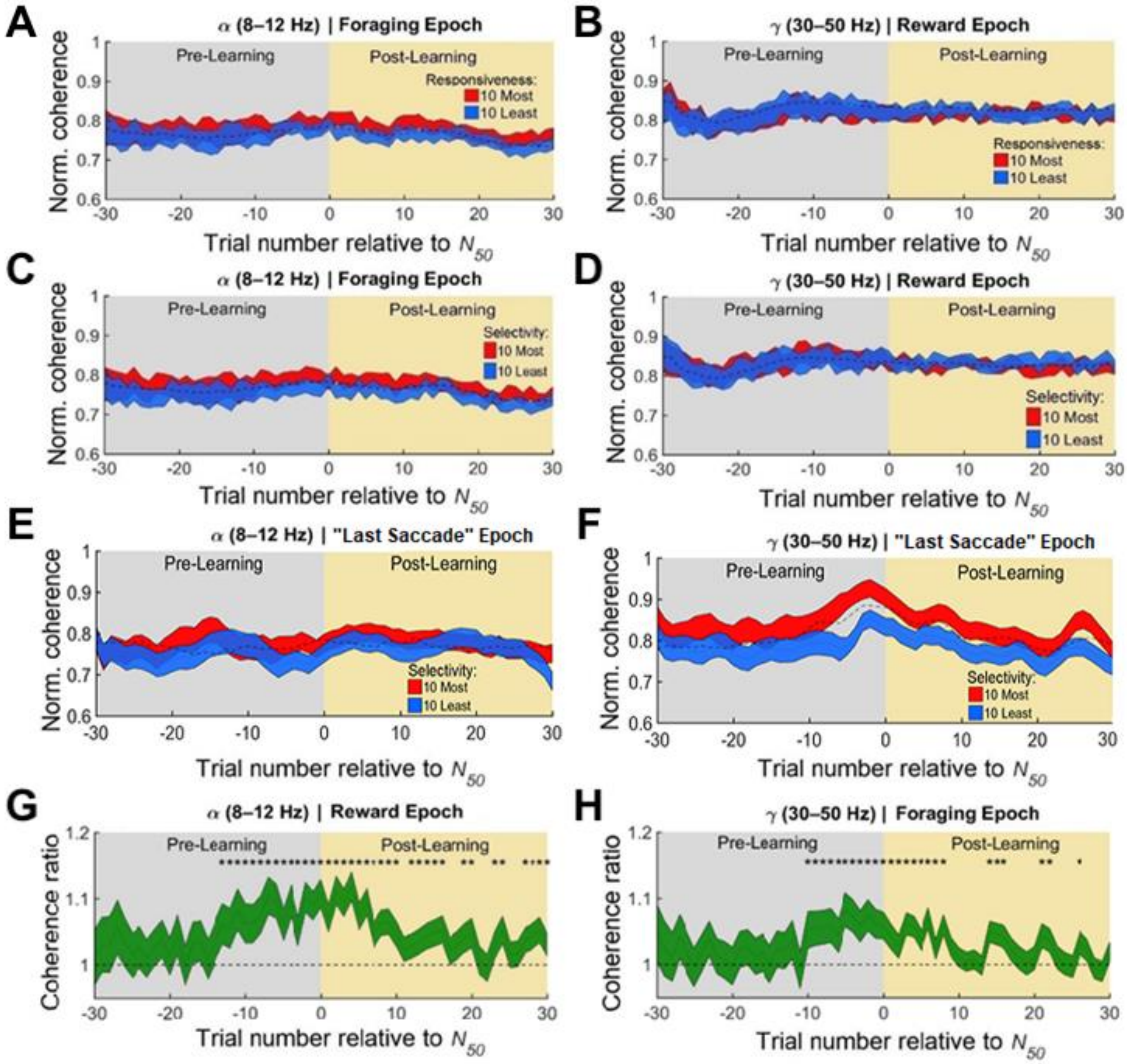

A
α (8–12 Hz) | Foraging Epoch
Pre-Learning
Post-Learning
Responsiveness:
10 Most
10 Least
Norm. coherence
Trial number relative to N50
B
γ (30–50 Hz) | Reward Epoch
Pre-Learning
Post-Learning
Responsiveness:
10 Most
10 Least
Norm. coherence
Trial number relative to N50
C
α (8–12 Hz) | Foraging Epoch
Pre-Learning
Post-Learning
Selectivity:
10 Most
10 Least
Norm. coherence
Trial number relative to N50
D
γ (30–50 Hz) | Reward Epoch
Pre-Learning
Post-Learning
Selectivity:
10 Most
10 Least
Norm. coherence
Trial number relative to N50
E
α (8–12 Hz) | "Last Saccade" Epoch
Pre-Learning
Post-Learning
Selectivity:
10 Most
10 Least
Norm. coherence
Trial number relative to N50
F
γ (30–50 Hz) | "Last Saccade" Epoch
Pre-Learning
Post-Learning
Selectivity:
10 Most
10 Least
Norm. coherence
Trial number relative to N50
G
α (8–12 Hz) | Reward Epoch
Pre-Learning
Post-Learning
Coherence ratio
Trial number relative to N50
H
γ (30–50 Hz) | Foraging Epoch
Pre-Learning
Post-Learning
Coherence ratio
Trial number relative to N50

#### Figure S3, related to Figure 4. Specificity of epochs and frequency bands.

(A-D) Normalized coherence between IT and PFC is shown for alpha (8-12 Hz) synchronization during foraging (left column) & low-gamma (30-50 Hz) synchronization following reward (right column). The frequency bands have been swapped across epochs relative to the data shown in Figure 4. Each band represents the grand median of all usable electrode pairs for all images. Error bars display standard error (SEM) across images.

(A,B) Average levels of synchronization, with the data split by *responsiveness* to the appearance of images in IT. For all epochs and bands, there was no statistical difference in the extent to which PFC was synchronized with more responsive (red) and less responsive (blue) sites in IT. The dashed black line indicates the overall average synchronization, which did not change noticeably across trials for these frequencies and these epochs.

(C,D) Average levels of synchronization, with the data split by electrode *selectivity* levels for images in IT. For alpha oscillations during the Foraging epoch and for gamma oscillations during the Reward epoch, there was no statistical difference in the extent to which PFC was synchronized with informative (red) and uninformative (blue) sites in IT. The dashed black line indicates the overall average synchronization, which did not change noticeably across trials for these frequencies and these epochs.

(E-F) Synchronization between IT and PFC is shown for the alpha band (E) and gamma band (F) in the "last saccade" epoch, which consists of the window preceding the saccade into the RZ (225ms to 100ms before the onset of the saccade) . These data shown in the same style as Figure 4. The most selective IT electrodes showed higher gamma synchronization (30-50 Hz) with PFC around the moment of learning in the Foraging epoch.

(G-H) Complementary figures for Figure 4A and Figure 4B:

(G) For Figure 4A, the ratio of synchronization strength for the most and least informative electrodes across trials relative to $N_{50}$. Asterisks indicate trials in which the ratio was significantly different from 1.

(H) For Figure 4B, as in (G), but for alpha synchronization (8-12 Hz) during the Reward epoch.

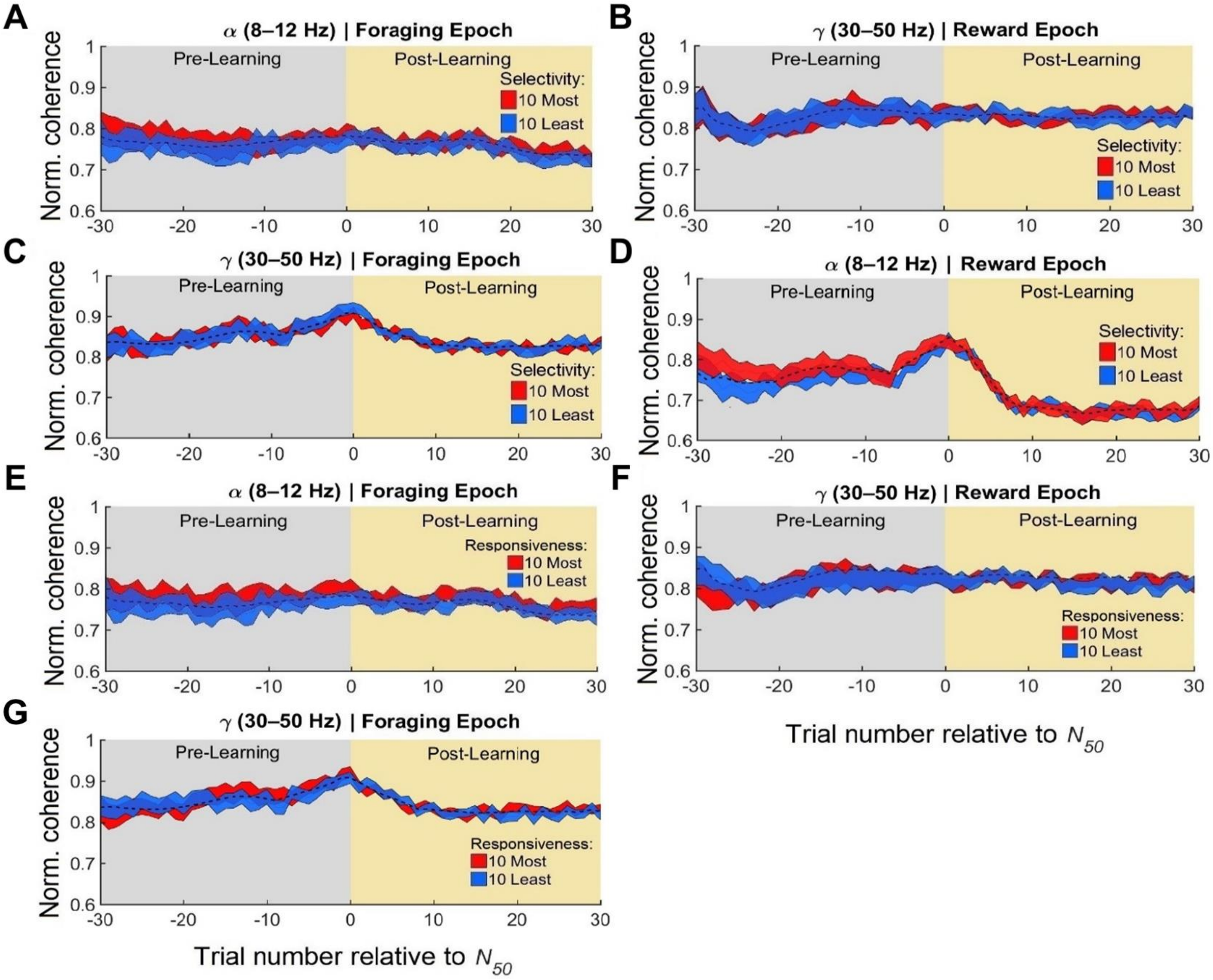
A
α (8–12 Hz) | Foraging Epoch
Pre-Learning
Post-Learning
Selectivity:
10 Most
10 Least
Norm. coherence
B
γ (30–50 Hz) | Reward Epoch
C
γ (30–50 Hz) | Foraging Epoch
D
α (8–12 Hz) | Reward Epoch
E
α (8–12 Hz) | Foraging Epoch
Responsiveness:
F
γ (30–50 Hz) | Reward Epoch
G
γ (30–50 Hz) | Foraging Epoch
Trial number relative to N50

#### Figure S4, related to Figure 5. Image responsive or selective sites in PFC do not unilaterally drive synchronization with IT.

Normalized coherence between IT and PFC is shown for during foraging (left column) and low-gamma synchronization following reward (right column) for alpha (8-12 Hz) and low-gamma (30-50 Hz) bands. Each band represents the grand median of all usable electrode pairs for all images. Error bars display standard error (SEM) across images.

(A-D) Average levels of synchronization, with the data split by electrode *selectivity* levels for images in PFC, as using PFC – rather than IT electrodes – to define selectivity. For all epochs and bands, there was no statistical difference in the extent to which IT was synchronized with informative (red) and uninformative (blue) sites in PFC. The dashed black line indicates the overall average synchronization, which did not change noticeably across trials for these frequencies and these epochs.

(E-G) Average levels of synchronization, with the data split by *responsiveness* to the appearance of images in PFC. For alpha oscillations during the Foraging epoch and for gamma oscillations during both the Foraging and the Reward epoch, there was no statistical difference in the extent to which IT was synchronized with more responsive (red) and less responsive (blue) sites in PFC. The dashed black line indicates the overall average synchronization, which did not change noticeably across trials for these frequencies and these epochs.

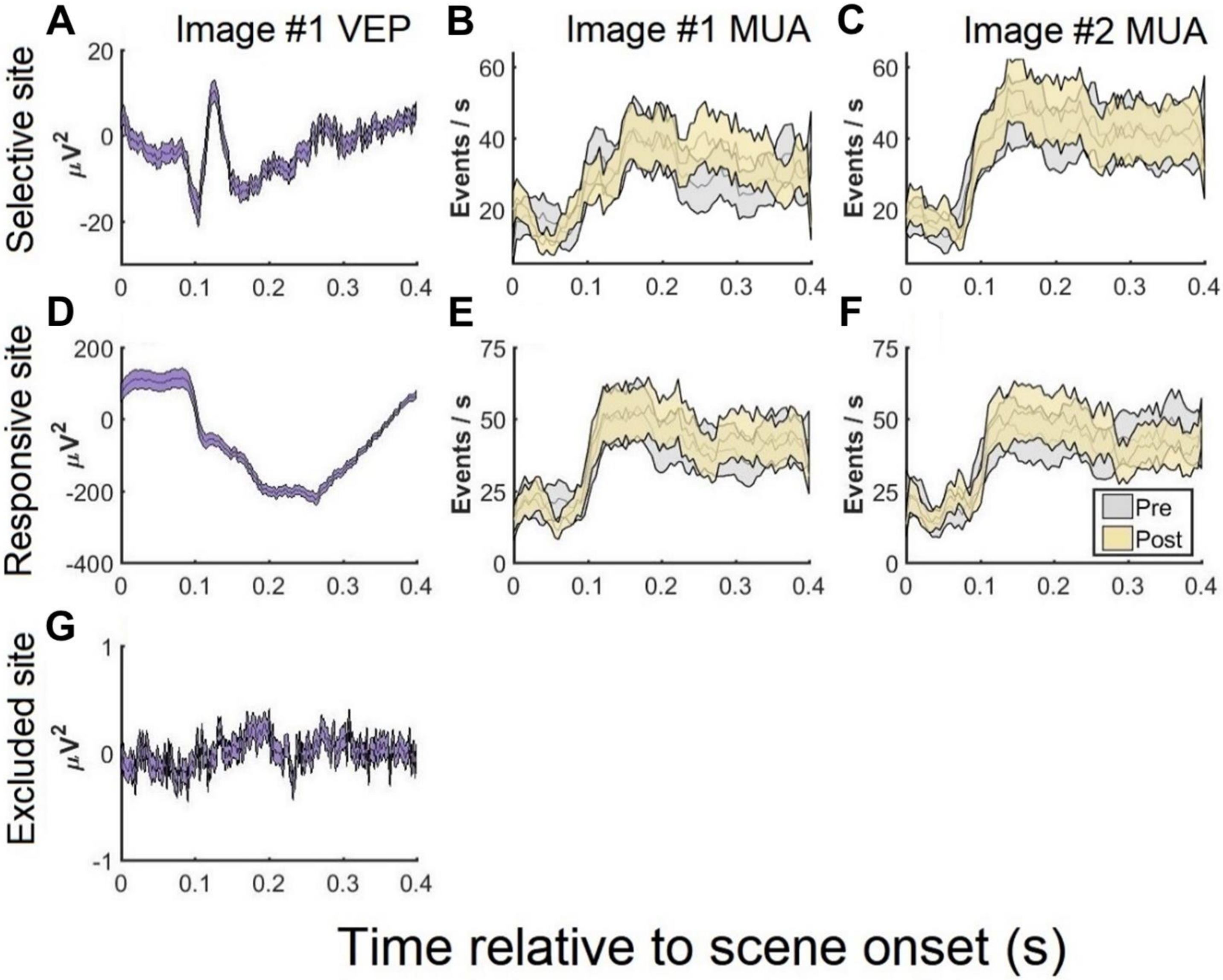
A
Image #1 VEP
B
Image #1 MUA
C
Image #2 MUA
Selective site
Events / s
D
E
F
Responsive site
Pre
Post
G
Excluded site
Time relative to scene onset (s)

**Figure S5, related to Figure 7. Examples of sensory responses in IT.**

(A) Visual-evoked potential from a selective IT channel is shown. The data presented are for the same session used in Figure 1.

(B,C) Multi-unit activity during scene onset is shown for the same channel used in (A) for two images from the same session. Yellow shows the data from the pre-learning phase of the session, while gray shows the data from the post-learning phase of the session. This site responds more to image 2 than to image 1, with no significant change in firing rate or selectivity across stages of learning.

(D) Visual-evoked potential from a more responsive and less selective IT channel are shown. This site responds strongly to image onset but does not distinguish between image 1 and image 2 and does not exhibit significant changes in responses across stages of learning.

(E,F) As in (B,C), but for (D).

(G) An example signal from an excluded IT channel is shown.

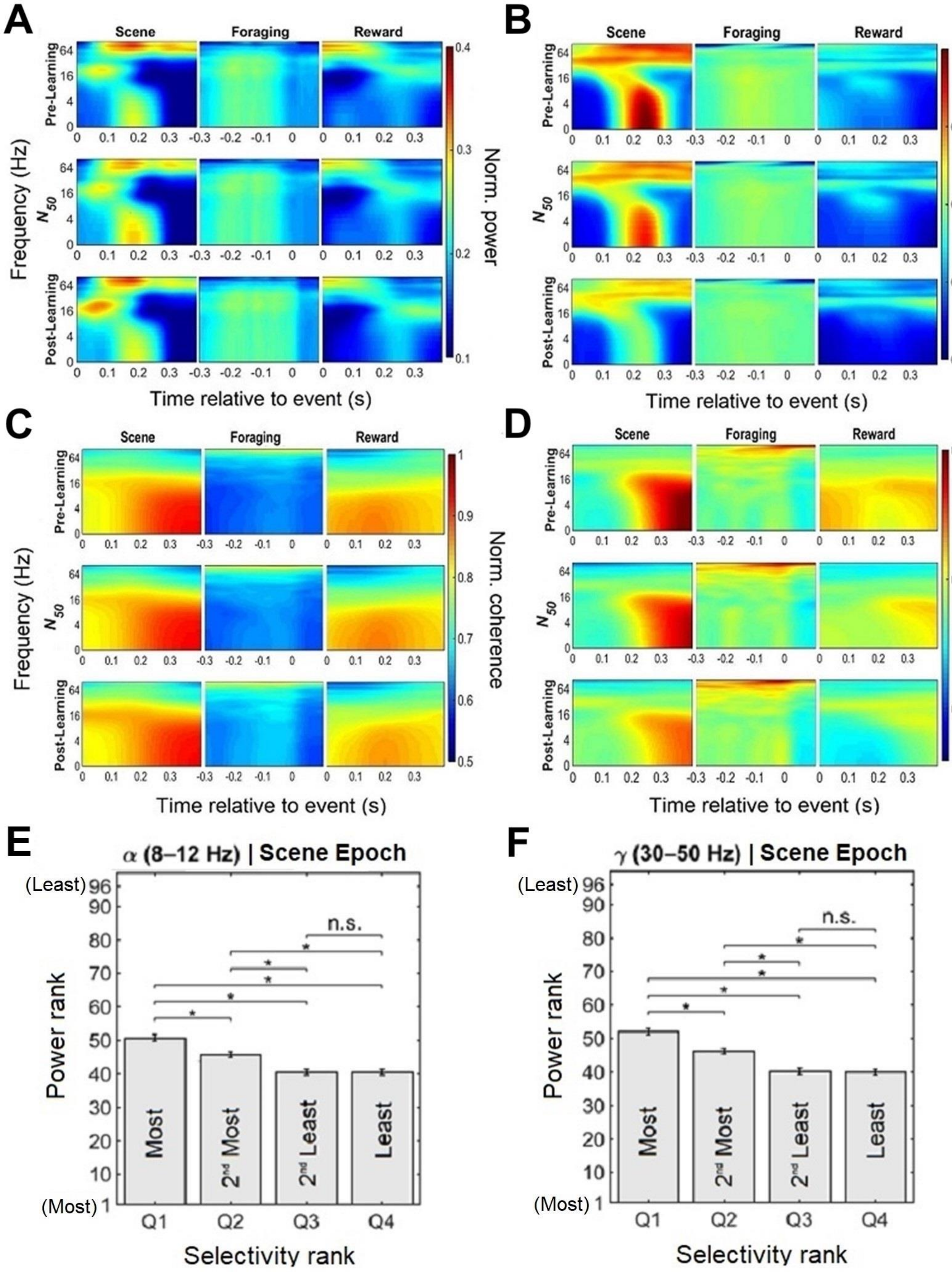
A
B
C
D
Scene
Foraging
Reward
Pre-Learning
N50
Post-Learning
Frequency (Hz)
Time relative to event (s)
Norm. power
Norm. coherence
E
α (8–12 Hz) | Scene Epoch
F
γ (30–50 Hz) | Scene Epoch
(Least)
(Most)
Power rank
Most
2nd Most
2nd Least
Least
n.s.
Q1
Q2
Q3
Q4
Selectivity rank

**Figure S6, related to Figure 7. Abrupt learning is not captured by local LFP power or synchronization.**

Time-frequency oscillatory power and coherence (1-100 Hz) for IT (A,C) and PFC (B,D) are shown for three trial epochs (‘Scene’, ’Foraging’, ’Reward’) and three stages of learning (‘Pre-Learning’, ’$N_{50}$’, ’Post-Learning). Learning epochs are binned and metrics are normalized as in Figure 3. Data shown are the grand average of all IT and PFC electrodes for all images for both animals. Reddish colors indicate high power or coherence, while bluish colors indicate low power or coherence.

(A) LFP power in area IT, across all frequencies, time points, trial epochs, and learning stages. The only difference across learning stages was a slight increase in beta (16-24 Hz) power after scene onset (lower left panel), which was confined to the time points earlier than visual response latencies (< 100 ms) and which did not reach statistical significance (two-way ANOVA, $p > 0.05$).

(B) LFP power in area PFC. There was a decrease in low-frequency power (4-12 Hz) during the Scene Onset epoch (two-way ANOVA, $p < 0.05$).

(C) LFP-LFP coherence within IT changed very little with learning.

(D) LFP-LFP coherence within PFC was largely unchanged with learning, except for a modest decrease in the low frequencies (1-12 Hz) from the pre-learning to the post-learning stage of the session (two-way ANOVA, $p < 0.05$).

(E-F) A relationship between IT sites’ image selectivity and power was tested for. Electrodes were ranked by their trial-averaged power (in each band individually) during the scene onset period. The results show that there is a very weak (negative) relationship between unit selectivity and power on the same electrodes. To visualize this relationship more clearly, we plot data from animal M, after dividing the selectivity values into quartiles.

(E) Ranking of trial-averaged alpha (8-12 Hz) power versus image selectivity ranking.

(F) Ranking of trial-averaged low-gamma (30-50 Hz) power during versus image selectivity ranking.

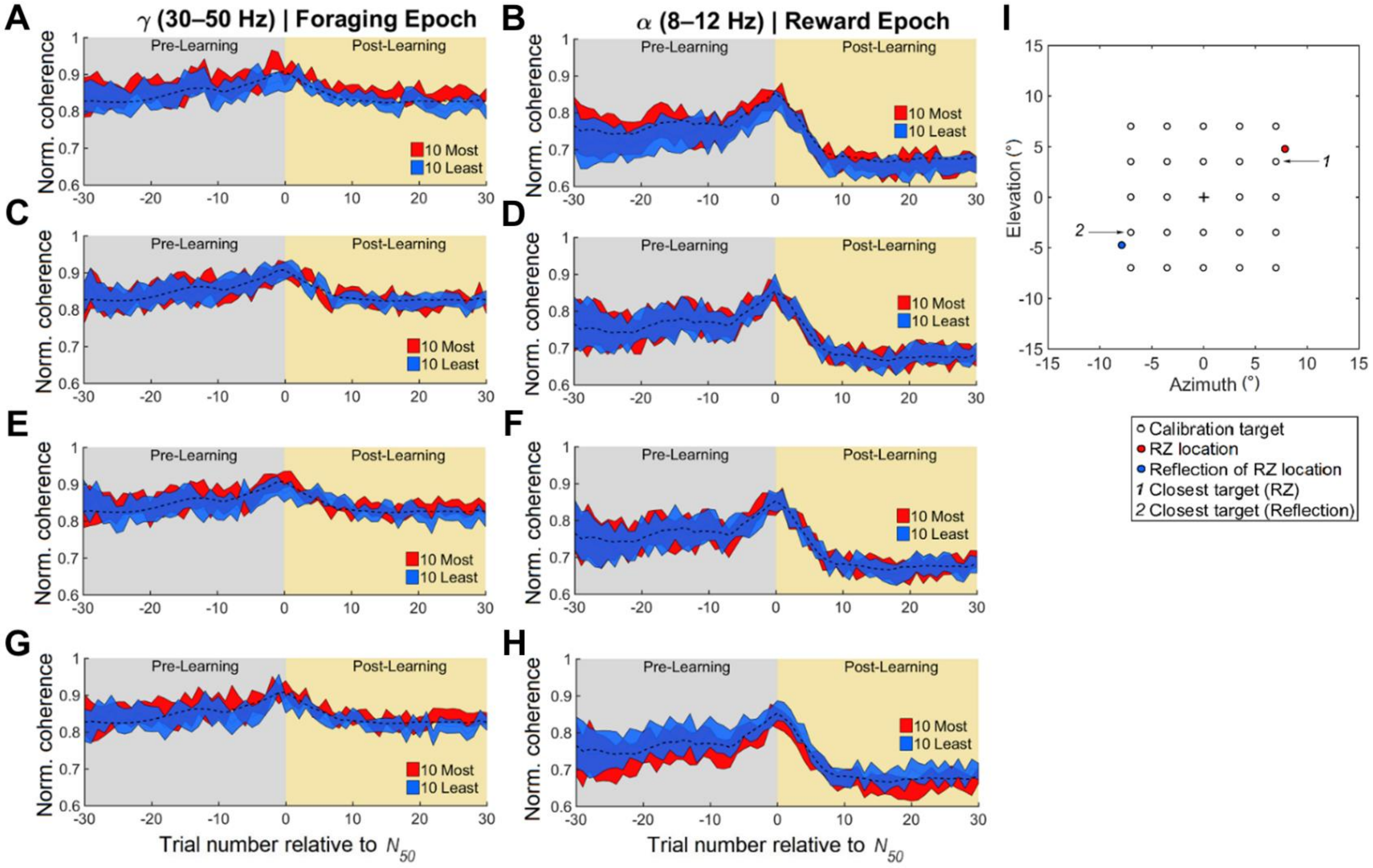
A
γ (30–50 Hz) | Foraging Epoch
B
α (8–12 Hz) | Reward Epoch
I
C
D
E
F
G
H
Pre-Learning
Post-Learning
Norm. coherence
10 Most
10 Least
Trial number relative to $N_{50}$
Elevation (°)
Azimuth (°)
1
2
Calibration target
RZ location
Reflection of RZ location
1 Closest target (RZ)
2 Closest target (Reflection)

#### Figure S7, related to Figure 5. Retinotopically selective or spatiotopically selective sites in PFC do not drive synchronization with IT, and vice-versa.

Normalized coherence between IT and PFC is shown for low-gamma (30-50 Hz) synchronization during foraging and alpha (8-12 Hz) synchronization following reward. Each band represents the grand median of all usable electrode pairs for all images. Error bars display standard error (SEM) across images. The dashed black line indicates the overall average synchronization, which did not change noticeably across trials for these frequencies and these epochs. For all conditions, there was no statistical difference in the extent to which informative (red) and uninformative (blue) sites in either PFC or IT synchronized with the other area.

(A,B) Average levels of synchronization, with the data split by electrode selectivity levels for *retinotopy* in PFC, as determined with a selectivity index using multi-unit activity on each electrode.

(C,D) As in (A,B), but with data split by electrode selectivity levels for *retinotopy* in IT, as determined with a selectivity index using multi-unit activity on each electrode.

(E,F) Average levels of synchronization, with the data split by electrode selectivity levels for *spatiotopy* in PFC, as determined with a selectivity index using multi-unit activity on each electrode.

(G,H) As in (E,F), but with data split by electrode selectivity levels for *spatiotopy* in IT, as determined with a selectivity index using multi-unit activity on each electrode.

(I) Diagram of target locations for the calibration task. On every trial, animals made a saccade to a small, high-contrast saccade target, randomly chosen from one of 9 (or 25; illustrated above) locations on a 3x3 (or 5x5; illustrated above) grid spanning the central 14 horizontal and vertical degrees on the monitor. To determine if MUAs encoded the spatiotopic location of the RZ, we matched RZ coordinates (red) and the reflection of the coordinates *(*blue) for each image to the closest target locations in the calibration task. To determine if MUA encoded the retinotopic location of the RZ, we sorted RZ locations according to their positions relative to initial fixation points on each foraging trial, and matched resulting vectors to saccades made in the same (or opposite) direction in the calibration task.
